# Activity-induced weakening of hippocampal input to prefrontal VIP interneurons reshapes inhibitory microcircuit function: implications for spatial working memory task acquisition

**DOI:** 10.1101/2025.07.21.665987

**Authors:** Shana E. Silverstein, Thomas T. Clarity, Meena S. Deshpande, Erik Vaughan, Shoshana Novik, Hector E. Yarur, Shiliang Zhang, Valerie S. Tsai, Rong Ye, Rachel M. Mikofsky, Madeline Hsiang, Avery Bauman, Gabriel Loewinger, Francisco Pereira, Marisela Morales, Vikaas S. Sohal, Hugo A. Tejeda, Joshua A. Gordon, David A. Kupferschmidt

## Abstract

Long-range projections from ventral hippocampus (vHPC) to medial prefrontal cortex (mPFC) support cognitive functions including spatial working memory (SWM). vHPC input targets multiple prefrontal interneuron classes, yet how hippocampal input recruits these populations *in vivo*, whether this recruitment is plastic, and how such plasticity influences cognition remain unknown. Here, we combined optical stimulation of mouse vHPC inputs with calcium recordings from discrete mPFC interneuron populations, revealing persistent activity-induced reweighting of hippocampal recruitment across prefrontal inhibitory microcircuits. *Ex vivo* electrophysiology and computational modeling implicated weakened monosynaptic hippocampal drive onto vasoactive intestinal polypeptide (VIP)-expressing interneurons in this circuit reconfiguration. Prior vHPC input stimulation and the schizophrenia-associated *Df(16)A^+/–^* mutation, which also yielded reduced hippocampal input onto VIP interneurons, produced convergent alterations in task-related VIP interneuron activity associated with poorer SWM task learning. These findings implicate hippocampal input to VIP interneurons as a key locus of plasticity capable of reconfiguring inhibitory microcircuit activity linked to impaired working memory task learning.

Graphical Abstract

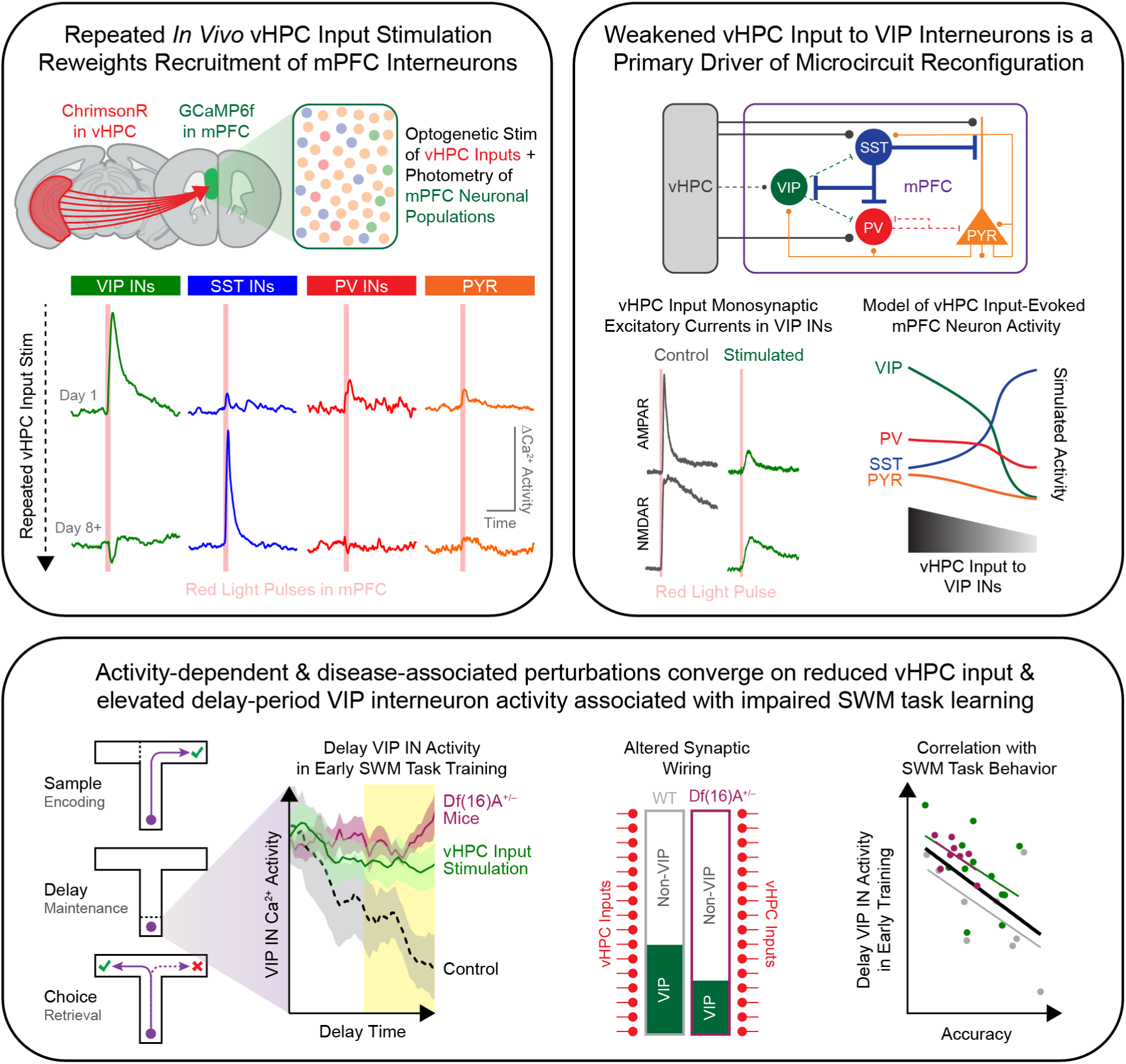

## INTRODUCTION

Functional interactions between the hippocampus and prefrontal cortex support adaptive behavior across diverse cognitive and emotional domains (Eichenbaum, 2017; Marek et al., 2018a; Shin and Jadhav, 2016; Sigurdsson and Duvarci, 2016; Zhang et al., 2021). By integrating information about past experience with ongoing behavioral demands, hippocampal-prefrontal interactions help guide contextually appropriate decisions and actions (Elston and Wallis, 2025; Shin and Jadhav, 2016; Yu and Frank, 2014). Reflecting their role in adaptive behavior, hippocampal-prefrontal interactions are dynamically regulated by experience (Padilla-Coreano et al., 2019; Parfitt et al., 2017; Tang et al., 2022), and their disruption has been linked to cognitive dysfunction in numerous neuropsychiatric disorders, including schizophrenia (Bähner and Meyer-Lindenberg, 2017; Cunniff et al., 2020; Godsil et al., 2013; Mukai et al., 2015; Shah et al., 2018; Sigurdsson et al., 2010).

In rodents, direct projections from ventral hippocampus (vHPC) to medial prefrontal cortex (mPFC) contribute to hippocampal-prefrontal oscillatory synchrony, influence prefrontal representations of behaviorally relevant information, and support learning and performance across multiple cognitive tasks, including spatial working memory (SWM) (Benchenane et al., 2010; Burette et al., 2000; Schoenfeld et al., 2023; Spellman et al., 2015; Yamamoto et al., 2014). Notably, vHPC inputs can induce persistent changes in vHPC-mPFC functional connectivity and cognitive function when repeatedly activated (Izaki et al., 2004; Laroche et al., 2000, 1990; Park et al., 2021; Ruggiero et al., 2021; Zheng and Zhang, 2015), suggesting that activity within this pathway can reshape prefrontal circuit function over extended timescales. Moreover, persistent alterations in vHPC input to mPFC have been linked to cognitive dysfunction in rodent models of neuropsychiatric disease (Cunniff et al., 2020; Mukai et al., 2015; Phillips et al., 2019; Sigurdsson et al., 2010; Tripathi et al., 2020). Together, these findings suggest that repeated vHPC activity can persistently reshape prefrontal circuit function. However, the mechanisms through which this occurs remain poorly understood.

vHPC axons provide direct excitatory input to multiple classes of mPFC interneurons (Ährlund-Richter et al., 2019; Canetta et al., 2020; Liu et al., 2020; Sun et al., 2019), positioning these populations to regulate how hippocampal signals influence prefrontal network function. Vasoactive intestinal polypeptide (VIP)-expressing interneurons preferentially inhibit other interneuron populations and thereby provide a powerful source of cortical disinhibition (Fu et al., 2014; Lee et al., 2013; Pi et al., 2013). In mPFC, VIP interneurons contribute to working memory maintenance, support task-related neural representations, and regulate the influence of hippocampal signals on prefrontal activity during behavior (Bae et al., 2025; Kamigaki and Dan, 2017; Lee et al., 2019). Another major target of vHPC input is the somatostatin (SST)-expressing interneuron population, which preferentially innervates pyramidal neuron dendrites and is well positioned to regulate integration of long-range excitatory inputs (Kepecs and Fishell, 2014; Markram et al., 2004; Rudy et al., 2011). Paralleling established roles for vHPC input, SST interneurons contribute to hippocampal-prefrontal synchrony, spatial information encoding, and SWM performance (Abbas et al., 2018). Altogether, these findings suggest that activity-dependent changes in how vHPC inputs recruit distinct interneuron populations may provide a mechanism by which experience persistently reshapes prefrontal microcircuit function. Yet, how hippocampal input recruits these populations *in vivo*, whether this recruitment exhibits activity-dependent plasticity, and how such plasticity influences cognition-related neural activity and behavior remain unclear.

The present study therefore asked whether repeated activation of vHPC inputs persistently alters recruitment of distinct prefrontal interneuron populations and how such plasticity influences circuit activity during cognition. To address this, we combined optogenetic stimulation of vHPC inputs with cell-type-specific recordings in mPFC. We found that vHPC inputs differentially recruit mPFC neuronal populations, with particularly robust engagement of VIP interneurons. Repeated stimulation of vHPC inputs produced cell-type-specific changes in neuronal recruitment, weakening VIP interneuron responses while enhancing SST interneuron activity. Slice electrophysiology and computational modeling identified reduced monosynaptic vHPC input onto VIP interneurons as a likely driver of these microcircuit changes. During early acquisition of a SWM task, mice with prior vHPC input stimulation exhibited elevated delay-period VIP interneuron activity that was associated with poorer task performance. Likewise, VIP interneurons in *Df(16)A^+/–^* mice, a model of schizophrenia-associated 22q11.2 deletion syndrome with impaired SWM task acquisition, exhibited elevated delay activity during early training and received proportionally fewer monosynaptic inputs from vHPC. Together, these findings identify activity-dependent reweighting of hippocampal recruitment across prefrontal inhibitory microcircuits and suggest that diminished hippocampal recruitment of VIP interneurons contributes to maladaptive prefrontal circuit activity associated with impaired working memory task learning.

## RESULTS

### Repeated activation of vHPC inputs to mPFC induces cell-type-specific plasticity in mPFC neuronal population recruitment

To characterize how vHPC input recruits distinct mPFC interneuron populations *in vivo*, we used an all-optical approach. The red-shifted opsin ChrimsonR was expressed in vHPC projection neurons and the Ca^2+^ indicator GCaMP6f in VIP or SST interneurons in mPFC. vHPC terminals were stimulated with red light while interneuron activity was measured simultaneously by fiber photometry through the same optical fiber implanted in mPFC (Figure 1A-C; S1-3).

**Figure 1.**
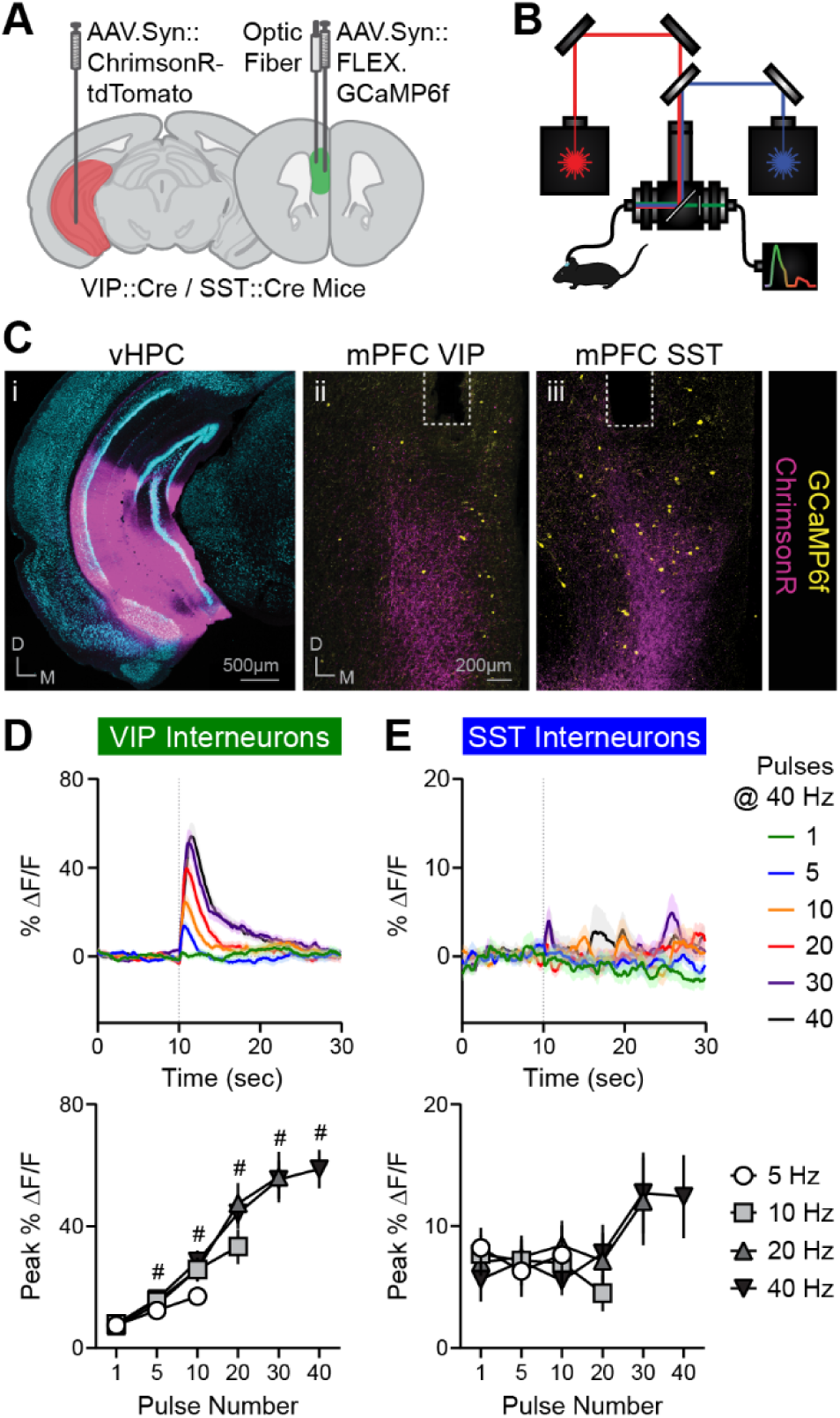
*In vivo* recruitment of mPFC VIP and SST interneurons by vHPC input. (**A**) Schematic showing AAV.Syn::ChrimsonR-tdTomato in unilateral vHPC, AAV.Syn::FLEX.GCaMP6f in unilateral mPFC, and an optic fiber in unilateral mPFC of VIP or SST::Cre mice. (**B**) Schematic of optogenetic stimulation-compatible, spectral-based photometry system. (**C**) Representative viral ChrimsonR-tdTomato expression (pink) in vHPC (i) and vHPC terminals in mPFC (ii,iii), GCaMP6f expression (yellow) in mPFC interneurons (VIP [ii] or SST [iii]) interneurons. Optic fiber placement denoted by dashed lines. DAPI staining (cyan) shown in vHPC image. (**D,E**) Top: Average vHPC input stimulation-evoked Ca^2+^ responses of mPFC VIP (**D**) and SST (**E**) interneurons in stimulation-naïve mice. Stimulation of different pulse numbers delivered at 40 Hz initiated at 10-sec timepoint. Bottom: Average peak vHPC input stimulation-evoked Ca^2+^ responses of VIP (**D**) and SST (**E**) interneurons in stimulation-naïve mice across a range of pulse numbers (1-40 pulses) and frequencies (5-40 Hz). n=23, 25 mice. VIP interneuron responses to 40-Hz stimulation: RM ANOVA, Main effect of Pulse Number: F(2.074, 45.64)=40.51, p<0.05; #p<0.05, different from 1 pulse.

Recruitment of VIP and SST interneurons by vHPC input was first characterized during an initial input-output (IO) session. During IO sessions, vHPC terminals in mPFC were stimulated using a range of pulse frequencies and train lengths (two rounds of 18 stimulation trains ranging from 1–40 pulses at 5–40 Hz; Figure S3B). VIP interneurons exhibited robust responses that scaled with increasing stimulation frequency and pulse number (Figure 1D; S4A), with maximal responses observed during the strongest stimulation condition (40 pulses at 40 Hz). In contrast, SST interneurons displayed comparatively weak responses that became detectable only at higher stimulation frequencies and pulse numbers (Figure 1E; S4G).

To determine whether recruitment of these interneuron populations by vHPC input exhibits activity-dependent plasticity, we developed a multi-day stimulation paradigm designed to assess both the induction and persistence of stimulation-induced changes in neuronal responses. Male and female mice underwent six input-output (IO) sessions across a 50-day period to longitudinally characterize responses to vHPC input stimulation (Figure 2A). A subset of these mice received an additional high-frequency stimulation (HFS) protocol to test whether repeated activation of vHPC inputs could drive persistent changes in interneuron response profiles. The HFS protocol consisted of 100 stimulation trains. Each train consisted of 40 pulses delivered at 40 Hz, corresponding to the IO condition that evoked the largest responses, with a 30-second inter-train interval to allow Ca²⁺ signals to return to baseline. This stimulation was administered over 12 days during the first three weeks of the experimental timeline, in addition to the six IO sessions (IO+HFS). The paradigm was informed by prior evidence linking vHPC input activity to hippocampal-prefrontal gamma oscillatory synchrony (Spellman et al., 2015), as well as pilot experiments demonstrating relative stability of neuronal responses within individual HFS sessions but pronounced changes across repeated stimulation sessions (Figure S3C).

**Figure 2.**
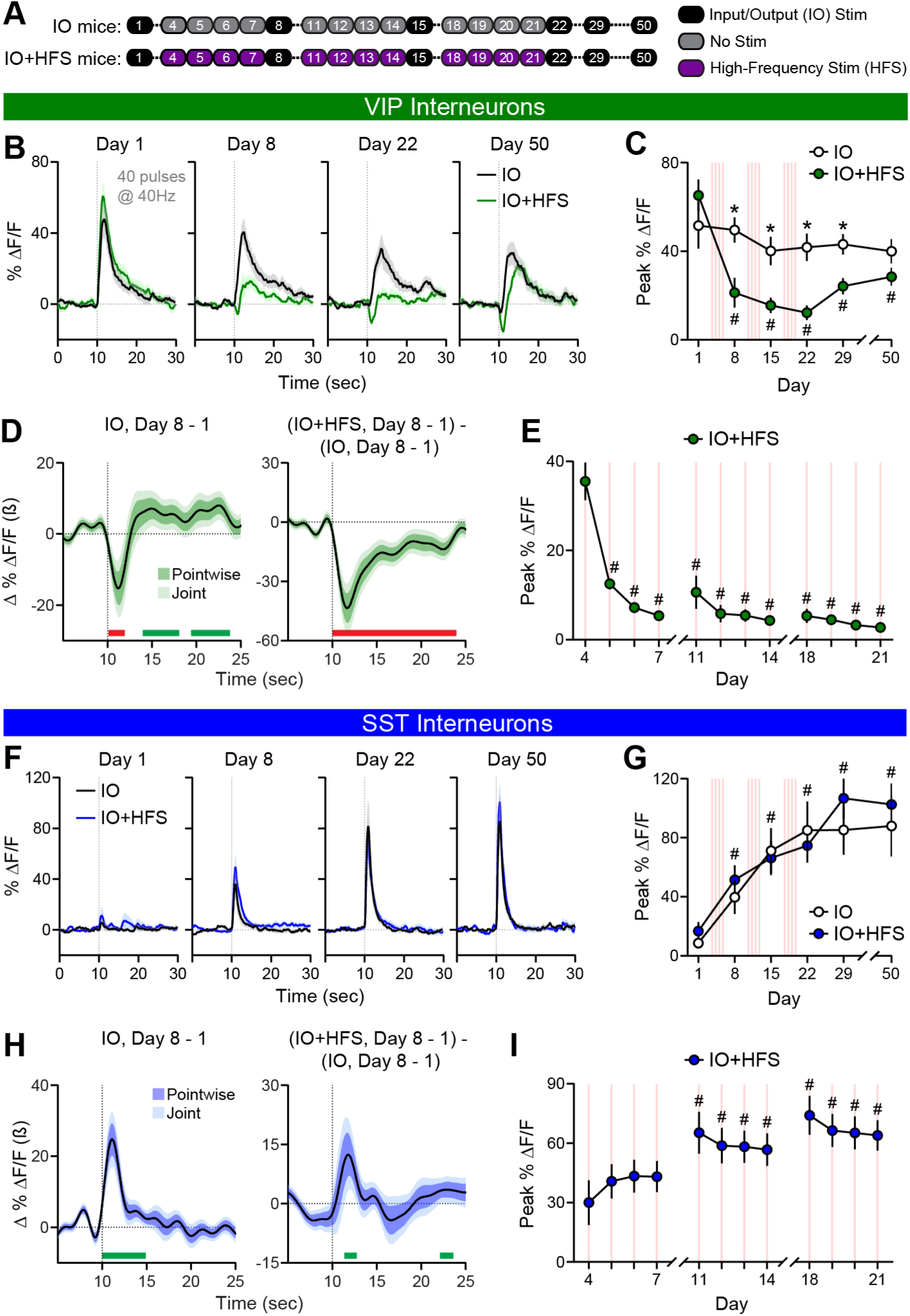
Repeated in vivo vHPC input stimulation induces persistent, cell-type-specific plasticity in recruitment of mPFC VIP and SST interneurons. (**A**) Schematic of 50-day experimental protocol for IO and IO+HFS mice. (**B**) Average stimulation-evoked Ca^2+^ responses of mPFC VIP interneurons in IO and IO+HFS mice on IO Days 1, 8, 22, and 50. Stimulation (40 pulses at 40 Hz) initiated at 10-sec timepoint. (**C**) Average peak stimulation-evoked Ca^2+^ responses of VIP interneurons in IO and IO+HFS mice across all six IO days. Stimulation was 40 pulses at 40 Hz. 2-way ANOVA, Main effect of Day: F(3.349, 70.33)=12.81, p<0.001; Main effect of Stimulation: F(1, 21)=7.903, p<0.05; Day x Stimulation interaction: F(5, 105)=6.416, p<0.0001; *p<0.005, different from HFS mice; #p<0.005 different from Day 1; n=11,12. (**D**) Functional linear mixed modeling (FLMM) of Day and HFS covariate effects on mouse-level VIP interneuron Ca^2+^ signals in IO and IO+HFS mice. Left: Functional coefficient estimates of the Day covariate at each timepoint in 30-sec stimulation trials, showing differences in average Ca^2+^ signals in Day 8 relative to Day 1 in IO mice. Right: Functional coefficient estimates of the interaction between the Day and HFS covariates, showing how changes in average Ca^2+^ signals from Day 1 to 8 in IO mice differ in IO+HFS mice. Green and red bars denote post-stimulation timepoints of significant enhancement and reduction, respectively. (**E**) Average peak stimulation-evoked Ca^2+^ responses of VIP interneurons in HFS mice across HFS days. RM ANOVA, Main effect of Day: F(2.498, 27.47)=14.56, p<0.0001; #p<0.05, different from Day 4; n=12. (**F**) Same as **B** but for SST interneurons. (**G**) Same as **C** but for SST interneurons. 2-way ANOVA, Main effect of Day: F(1.934, 44.49)=29.48, p<0.0001; #p<0.001, different from Day 1; n=12,13. (**H**) Same as **D** but for SST interneuron Ca^2+^ signals. (**I**) Same as **E** but for SST interneurons. RM ANOVA, Main effect of Day: F(1.998, 21.97)=16.88, p<0.0001; #p<0.05, different from Day 4; n=12.

VIP interneuron responses to vHPC input stimulation gradually declined across IO sessions in mice receiving IO stimulation alone (Figure 2B,C; S4A,E). Modest reductions in peak response amplitude across IO days (Figure 2B,C; S4A,E) were accompanied by more pronounced reductions in activity during the stimulation period and delayed response kinetics (Figure S4A-D). In IO+HFS mice, VIP interneuron responses to vHPC input stimulation were rapidly and robustly attenuated (Figure 2B-D; S4E). This attenuation persisted across subsequent IO sessions, with partial recovery observed in the weeks following the final HFS session (Figure 2B,C). Importantly, attenuation of VIP interneuron recruitment was evident across a broad range of stimulation pulse numbers and frequencies in both IO and IO+HFS mice (Figure S4E). Consistent with these findings, VIP interneuron responses to HFS progressively declined across the first several HFS sessions (Figure 2E) and within the initial HFS session itself (Figure S4F). Together, these findings indicate that repeated recruitment of VIP interneurons by vHPC input results in progressive and persistent depression of the efficacy of this input.

In striking contrast, SST interneuron responses to vHPC input stimulation during IO sessions gradually increased over the 50-day experimental period (Figure 2F,G; S4G). This potentiation persisted throughout a three-week stimulation-free interval (Days 29-50) and was evident across multiple stimulation pulse numbers and frequencies (Figure S4G). IO+HFS mice showed a modest early enhancement of SST interneuron recruitment by vHPC input relative to IO mice (Figure 2H). However, the overall magnitude of potentiation remained largely comparable between IO and IO+HFS mice (Figure 2F-H). SST interneuron responses to HFS likewise increased across repeated HFS sessions (Figure 2I) but were unchanged within the initial HFS session (Figure S4H). Thus, repeated activation of the same hippocampal projection produced opposing plastic responses in VIP and SST mPFC interneurons.

These results indicate that repeated activation of vHPC inputs during IO stimulation alone was sufficient to induce plasticity in VIP and SST interneuron responses, and these adaptations were generally enhanced by additional HFS. To determine whether repeated HFS could independently induce these adaptations, we administered HFS without intervening IO sessions in a separate cohort of mice. HFS alone recapitulated the plasticity, yielding marked attenuation and enhancement of VIP and SST interneuron responses, respectively (Figure S5). Together, these findings demonstrate that repeated activation of vHPC inputs is sufficient to drive bidirectional, cell-type-specific changes in interneuron recruitment.

To place the cell type-specific plasticity observed in VIP and SST interneurons within a broader microcircuit context, we also recorded responses from parvalbumin (PV)-expressing interneurons and CaMKIIα-expressing putative pyramidal neurons. Like VIP interneurons, PV interneuron recruitment progressively declined with repeated vHPC input stimulation (Figure S4A-C). CaMKIIα-expressing putative pyramidal neurons showed a modest attenuation of evoked responses following repeated vHPC input stimulation, particularly during the post-stimulation period (Figure S6D-F). No sex differences were detected in any recorded cell population. Together, these findings demonstrate that repeated activation of vHPC inputs induces persistent, cell-type-specific plasticity across mPFC neuronal populations, with the most pronounced adaptations occurring among inhibitory interneuron classes.

### Prior vHPC input stimulation reduces vHPC monosynaptic connectivity with mPFC VIP interneurons

The pronounced, bidirectional changes observed in *in vivo* VIP and SST interneuron recruitment raised the possibility that repeated vHPC activation alters monosynaptic hippocampal drive onto these populations. To test this hypothesis, we assessed monosynaptic vHPC connectivity onto mPFC VIP and SST interneurons using whole-cell electrophysiology. Because HFS alone was sufficient to induce robust plasticity *in vivo*, mice received either 12 HFS sessions or no stimulation (NS) prior to *ex vivo* whole-cell electrophysiology recordings (Figure 3A-C). One day after the final stimulation session, optogenetically evoked excitatory postsynaptic currents (oEPSCs) were recorded from identified VIP and SST interneurons in acute mPFC slices. Monosynaptic currents were isolated using TTX and 4-AP and evoked by brief photostimulation of ChrimsonR-expressing vHPC terminals (Figure 3D). AMPA receptor (AMPAR)- and NMDA receptor (NMDAR)-mediated currents were measured separately using standard biophysical and pharmacological approaches (Linders et al., 2022; Petreanu et al., 2009) (Figure 3E; S7A,C).

**Figure 3.**
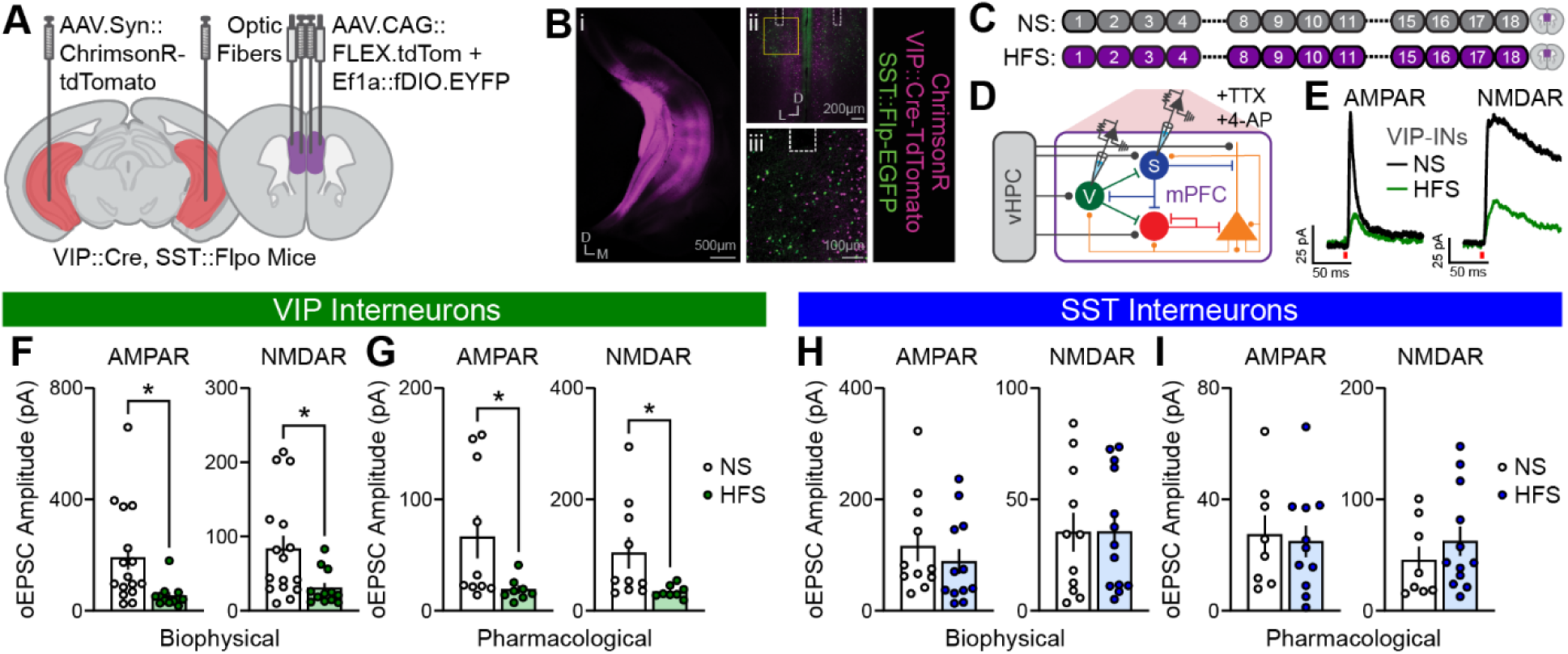
Prior vHPC input stimulation weakens monosynaptic vHPC input onto mPFC VIP interneurons. (**A**) Schematic showing AAV.Syn::ChrimsonR-tdTomato in bilateral vHPC, AAV:CAG::FLEX.tdTomato and AAV:Ef1a::fDIO.EYFP in bilateral mPFC, and optic fibers in bilateral mPFC of VIP::Cre, SST::Flpo mice. (**B**) Representative viral ChrimsonR-tdTomato expression (magenta) in vHPC (i) and vHPC terminals in mPFC (ii-iii), tdTomato expression (magenta) in VIP interneurons and EGFP expression (green) in SST interneurons (ii-iii). Optic fiber placement denoted by dashed lines. (**C**) Schematic of 18-day experimental protocol consisting of either 12 HFS sessions or no stimulation (NS). Euthanasia and recordings were conducted 24 hours following final stimulation. (**D**) Schematic of brain slice whole-cell electrophysiology recording of monosynaptic AMPA receptor (AMPAR)- and NMDA receptor (NMDAR)-mediated currents in VIP or SST interneurons evoked by optogenetic stimulation of vHPC inputs to mPFC. (**E**) Representative vHPC input stimulation-evoked monosynaptic AMPAR- and NMDAR-mediated currents (pharmacologically isolated) in VIP interneurons of a NS or HFS mouse. (**F**) Average biophysically isolated AMPAR- and NMDAR-mediated monosynaptic oEPSC amplitude in VIP interneurons of NS and HFS mice. AMPAR: *p<0.005, U=33; NMDAR: *p<0.01, U=40; Mann Whitney tests; n=16,12 cells, 13,10 mice. (**G**) Average pharmacologically isolated AMPAR- and NMDAR-mediated monosynaptic oEPSC amplitude in VIP interneurons of NS and HFS mice. Mann-Whitney tests, AMPAR: *p<0.05, U=15; NMDAR: *p<0.01, U=11; n=10,8 cells, 9,6 mice. (**H**) Average biophysically isolated AMPAR- and NMDAR-mediated optically evoked monosynaptic oEPSC amplitude in SST interneurons of NS and HFS mice; n=11,13 cells, 9,11 mice. (**I**) Average pharmacologically isolated AMPAR- and NMDAR-mediated monosynaptic oEPSC amplitude in SST interneurons of NS and HFS mice; n=8,12 cells, 7,11 mice.

Consistent with a weakening of vHPC recruitment of VIP interneurons observed *in vivo*, both AMPAR- and NMDAR-mediated oEPSCs were markedly reduced in VIP interneurons from HFS mice relative to NS controls (Figure 3F,G), with no accompanying change in AMPAR/NMDAR ratios (Figure S7B). In contrast, neither AMPAR- nor NMDAR-mediated currents were altered in SST interneurons from the same HFS and NS animals (Figure 3H,I); AMPAR/NMDAR ratios likewise were unchanged in SST interneurons (Figure S7D). These findings identify a selective reduction in monosynaptic excitatory drive onto VIP interneurons as a potential synaptic mechanism underlying the cell-type-specific plasticity induced by repeated vHPC activation.

### Computational modeling identifies reduced vHPC recruitment of VIP interneurons as a plausible driver of microcircuit plasticity

The selective weakening of monosynaptic vHPC drive onto VIP interneurons raised the possibility that this adaptation could contribute to the broader pattern of activity-dependent plasticity observed *in vivo*. Specifically, reduced recruitment of VIP interneurons by vHPC input may disinhibit SST interneurons, leading to enhanced SST responses despite unchanged monosynaptic vHPC drive. Increased SST interneuron activity could, in turn, suppress PV interneuron activity and further constrain VIP interneuron responses. The increase in SST interneuron-mediated inhibition on pyramidal neurons may outweigh the reduced PV interneuron-mediated inhibition to produce a modest attenuation of pyramidal neuron responses (Figure 4A).

**Figure 4.**
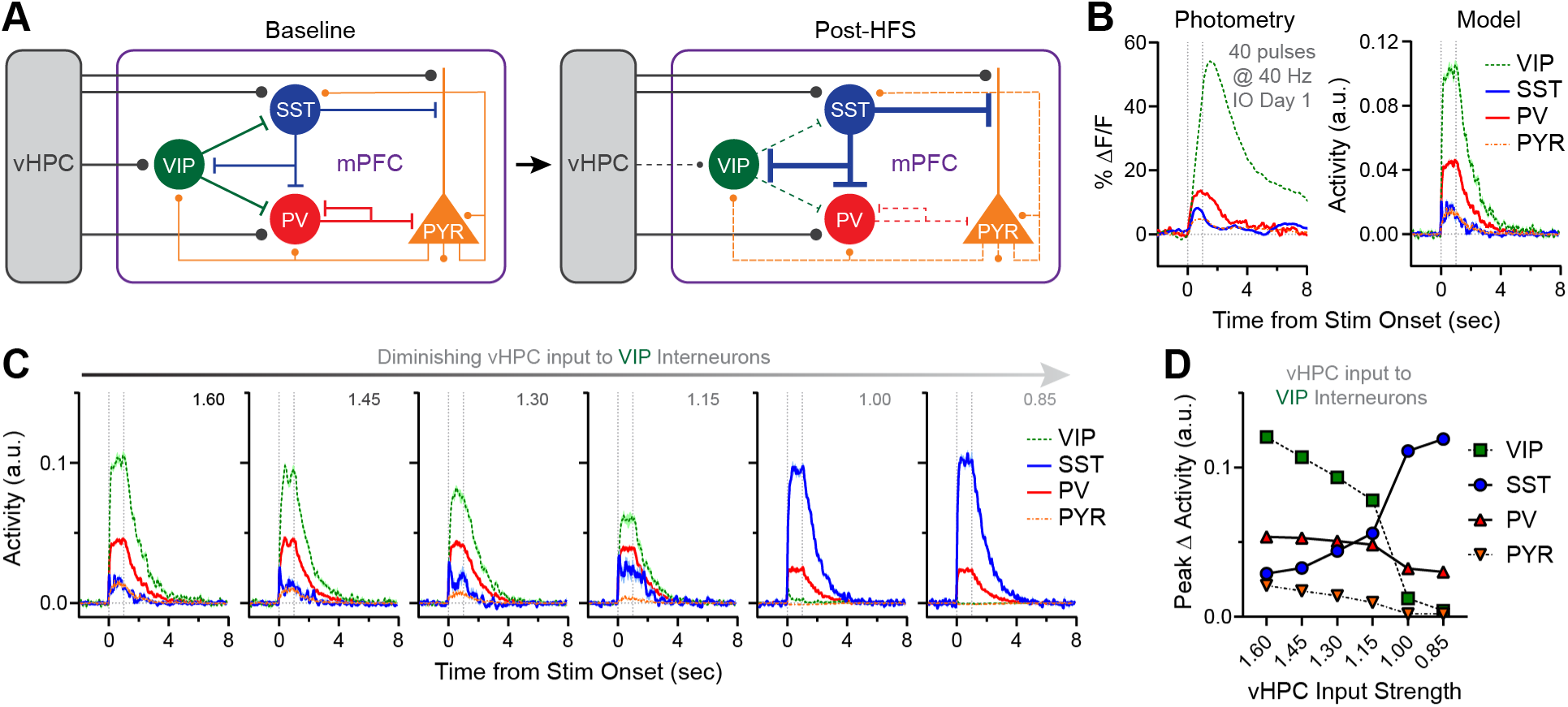
Computational modeling suggests that weakened vHPC input onto VIP interneurons is sufficient to recapitulate observed microcircuit adaptations. (**A**) Conceptual model of hypothesized changes in mPFC microcircuit following vHPC input HFS. (**B**) Comparison of Ca^2+^ activity (left) and computationally modeled activity (right) in response to real or modeled vHPC input stimulation in mPFC neuronal populations or their model analogs. Left: Average stimulation-evoked Ca^2+^ responses of mPFC neuronal populations on IO Day 1. Right: Simulated activity from integrate-and-fire computational model of mPFC microcircuit evoked by stimulation of “vHPC” input. (**C**-**D**) Simulated traces (**C**) and peak activity (**D**) for each neuronal population during responses to vHPC input stimulation, shown for diminishing strengths of vHPC input to VIP interneurons (values ranging from 1.60 [initial] to 0.85).

To evaluate whether reduced vHPC input onto VIP interneurons could plausibly account for these population-level changes, we developed a computational model incorporating the four principal neuronal classes examined experimentally. The model consisted of 36 integrate-and-fire neurons, including VIP (“disinhibitory”), SST (“feedback”), PV (“feedforward”), and pyramidal (PYR) neurons, connected according to canonical patterns of cortical microcircuit connectivity informed by experimental studies (Canetta et al., 2020; Petreanu et al., 2009; Pfeffer et al., 2013) (Figure S8). All neuron classes received excitatory drive from an external source representing vHPC input. During each 10-second simulation, vHPC input was delivered simultaneously to all neuronal populations for 1 second. Model parameters were modestly adjusted to yield stable network dynamics and qualitatively reproduce the relative response magnitudes observed experimentally during the initial IO session (40 pulses at 40 Hz; Figure 4B). We then systematically varied the strength of vHPC input onto VIP interneurons while holding all other parameters constant.

As expected, reducing vHPC input onto VIP interneurons progressively decreased evoked VIP interneuron activity in the model (Figure 4C,D). Importantly, this manipulation also produced a marked enhancement of SST interneuron activity. Consistent with our *in vivo* photometry data, PV interneuron activity and pyramidal neuron activity likewise declined as vHPC input onto VIP interneurons was reduced (Figure S6). Thus, canonical patterns of cortical microcircuit connectivity were sufficient to reproduce the qualitative pattern of cell-type-specific plasticity observed experimentally based solely on a reduction of vHPC input onto VIP interneurons. Together with the slice electrophysiology findings, these results identify reduced monosynaptic vHPC input onto VIP interneurons as a plausible circuit mechanism capable of driving the broader microcircuit plasticity induced by repeated vHPC activation.

### Activity-dependent microcircuit plasticity occurs without significant impairment in SWM acquisition

To examine how activity-dependent plasticity in hippocampal-prefrontal inhibitory circuits influences SWM-related neural activity and behavior, mice expressing ChrimsonR in vHPC and GCaMP6f in either VIP or SST interneurons received either 12 sessions of HFS or NS prior to training on a delayed non-match-to-sample (DNMS) T-maze task (Figure 5A-C; S9, S10A). These were the same mice in which HFS produced robust attenuation of VIP interneuron recruitment and potentiation of SST interneuron recruitment by vHPC input (Figure S5).

**Figure 5.**
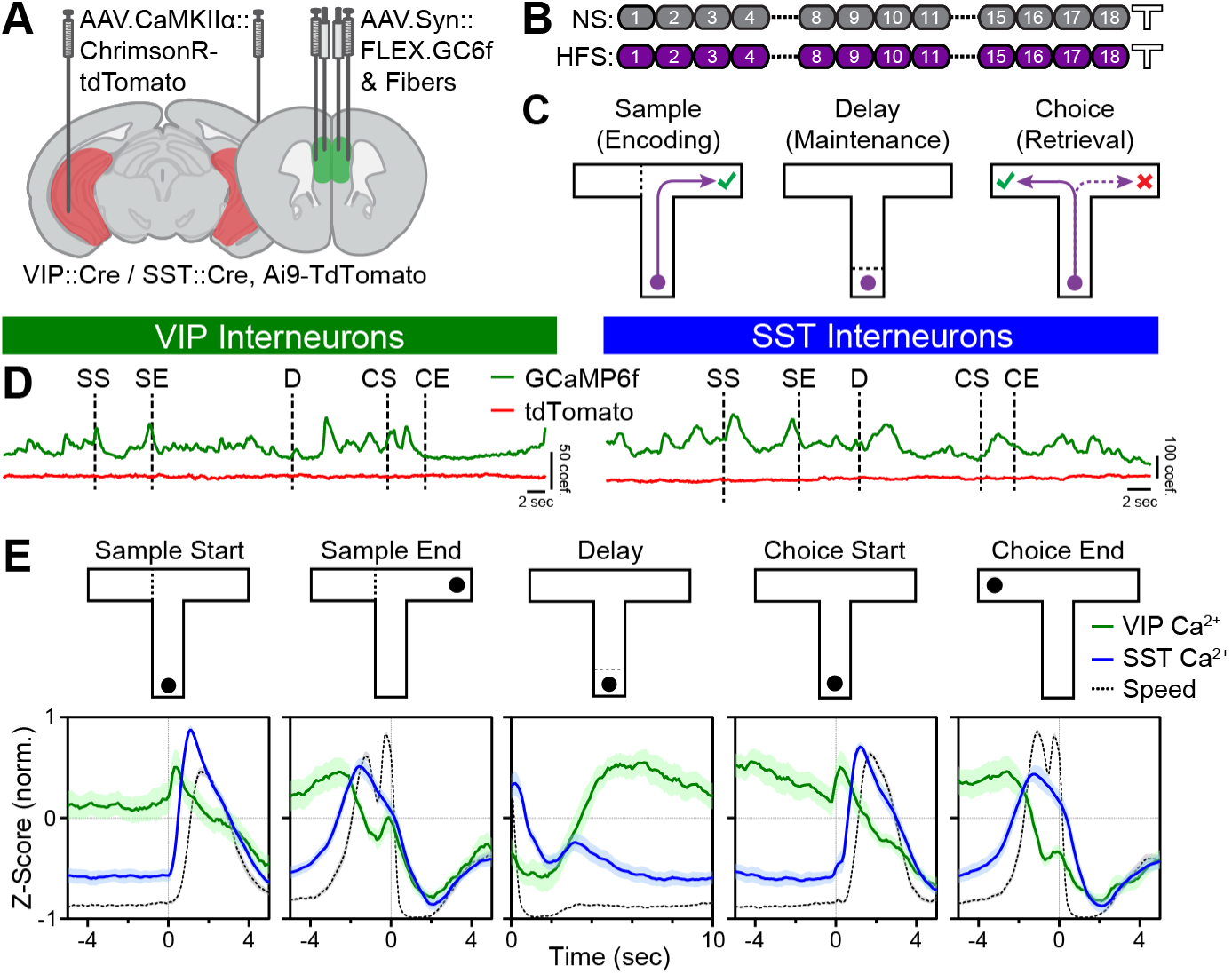
Distinct activity dynamics of mPFC VIP and SST interneurons across SWM task periods. (**A**) Schematic showing AAV.CaMKIIα::ChrimsonR-tdTomato in bilateral vHPC, AAV.Syn::FLEX.GCaMP6f in bilateral mPFC, and optic fibers in bilateral mPFC of VIP or SST::Cre, Ai9-tdTomato mice. (**B**) Schematic of 18-day stimulation protocol consisting of either 12 sessions of no stimulation (NS) or HFS followed by training on SWM task. (**C**) Schematic of DNMS T-maze task. (**D**) Representative photometry traces of GCaMP6f and tdTomato fluorescence (after spectral unmixing) across a single trial from VIP and SST interneuron populations. Task events denoted by dashed lines: Sample Start (SS); Sample End (SE); Delay (D); Choice Start (CS); Choice End (CE). (**E**) Average normalized Ca^2+^ activity of VIP (green) and SST interneuron populations (blue), and average normalized mouse speed (gray dashed, combined VIP and SST interneuron mice) aligned to discrete task events of the SWM task. Data are normalized such that Z-scored Ca^2+^ signals from individual trials ranged from –1 to 1. Data are from correct trials performed by NS mice only. n=10-14 for Ca^2+^, n=24 for speed.

Following the final HFS or NS session, mice were trained on a DNMS task using a 10-second delay period (Hupalo et al., 2026; Passecker et al., 2025; Spellman et al., 2015). HFS mice acquired the task at a rate comparable to NS controls, despite a non-significant rightward shift in the distribution of days required to reach performance criterion (Figure S10B,C). Similarly, overall training accuracy did not differ between groups (Figure S10D). No sex differences were detected in behavioral performance. Despite robust activity-dependent remodeling of vHPC recruitment of VIP and SST interneurons, repeated vHPC activation did not significantly impair SWM task acquisition. We therefore next examined whether prior vHPC stimulation altered task-related interneuron activity during SWM learning.

### VIP and SST interneurons exhibit distinct activity dynamics during SWM task performance

Using photometry data from all training days and mice, Ca^2+^ activity was derived using a validated spectral unmixing procedure (Brockway et al., 2023; Meng et al., 2018; Sciolino et al., 2022) and aligned to five key task events: Sample Start, Sample End, Delay, Choice Start, and Choice End (Figure 5D,E; S11A-D). VIP and SST interneurons exhibited both aligned and distinct average activity profiles across SWM task events. Both populations showed similar patterns of task-related modulation during sample and choice periods, with increases at the start of these periods and decreases around goal approach and reward consumption; however, they differed in their temporal profiles. During the delay period, average VIP interneuron activity gradually increased whereas SST interneuron activity progressively decreased (Figure 5E; S12A,B).

Because VIP and SST interneuron activity was correlated with movement speed during task performance (Figure 5E, S12C), training-related changes in activity during task periods accompanied by movement could reflect alterations in locomotor behavior rather than neural processes associated with task learning. Indeed, movement speed changed substantially across training during the sample and choice periods but remained stable throughout the delay period (Figure S13). We therefore focused subsequent analyses on delay-period interneuron activity to identify neural dynamics associated with successful SWM task acquisition.

### Distal delay VIP interneuron activity predicts SWM task acquisition

To examine whether delay-period VIP interneuron activity relates to trial outcome, and how this relationship evolves across SWM training, we compared activity during correct and incorrect trials. During early training, VIP interneurons showed a decrease in activity during the final portion of the delay period (“distal delay”) on correct trials (Figure 6A,B). This decrease was smaller during incorrect trials and was absent during late stages of training. Indeed, distal delay VIP interneuron activity was elevated in incorrect relative to correct trials early in training, increased across training during correct trials, and increased less prominently during incorrect trials (Figure 6B; S15A). Collectively, these findings indicate that VIP interneuron activity during the distal delay period distinguishes subsequent correct and incorrect choices specifically during the early stages of SWM task acquisition. This relationship was not observed in SST interneurons, whose distal delay activity did not differ between correct and incorrect trials across training (Figure S15B). Notably, VIP and SST activity both differed between correct and incorrect trials during choice reward consumption (Figure S12); this activity did not differ by stimulation condition or training phase (data not shown).

**Figure 6.**
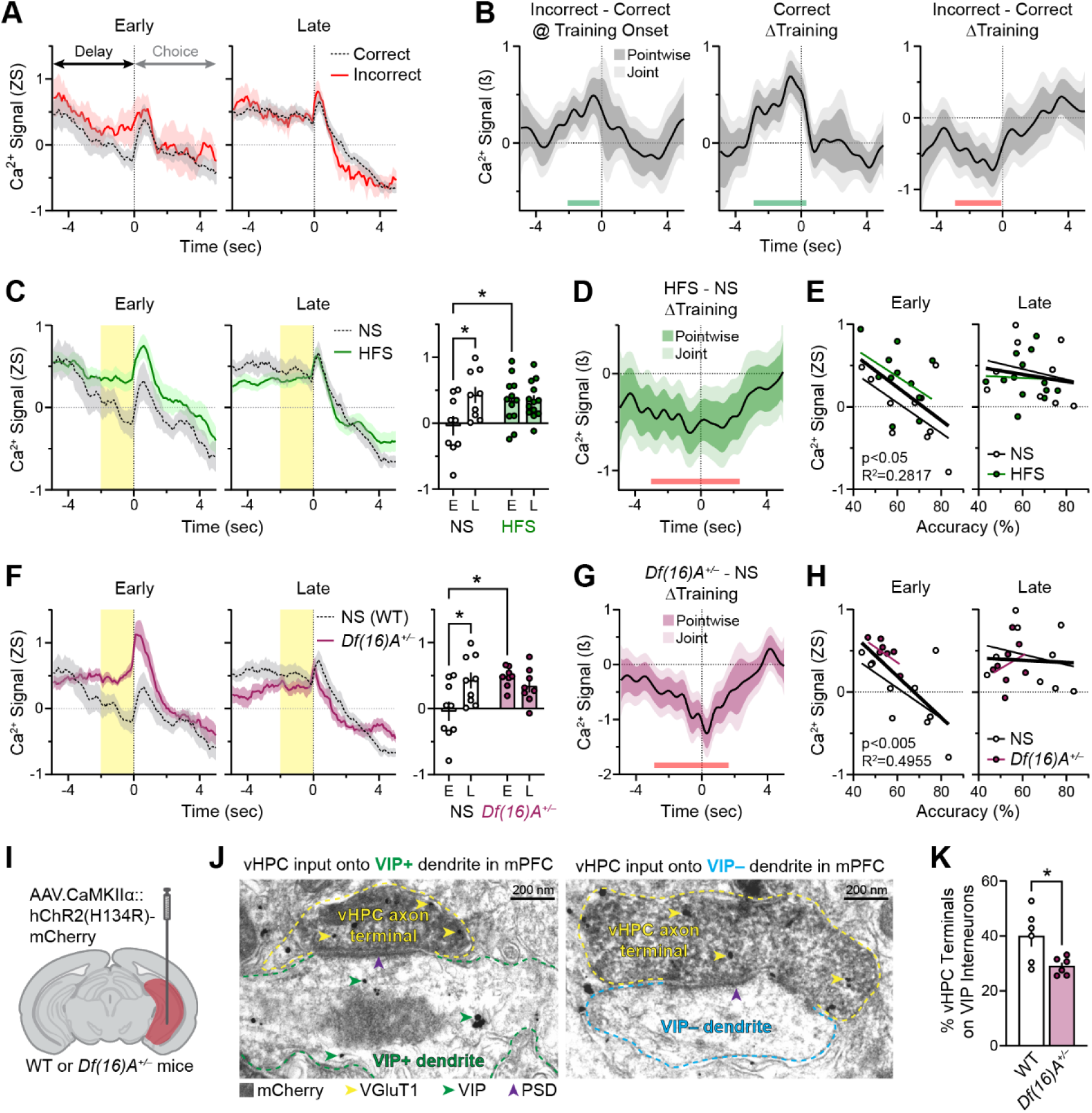
Activity-dependent and disease-associated perturbations converge on reduced vHPC input to VIP interneurons and elevated delay-period VIP interneuron activity associated with impaired SWM task learning. (**A**) Average Z-scored Ca^2+^ signals from VIP interneuron populations during the Delay-to-Choice transition for correct and incorrect trials in NS mice in early (first third of trials) and late (final third of trials) stages of training (n=7-9). (**B**) Functional linear mixed modeling (FLMM) of Outcome and Training covariate effects on trial-level VIP interneuron Ca^2+^ signals in NS mice during the Delay-to-Choice transition. Left: Functional coefficient estimates of the Outcome covariate at each timepoint in incorrect trials, showing differences in average Ca^2+^ signals in incorrect relative to correct trials at the onset of training. Middle: Functional coefficient estimates of the Training covariate in correct trials, showing changes in average Ca^2+^ signals across training. Right: Functional coefficient estimates of the interaction between the Training and Outcome covariates, showing how changes in average Ca^2+^ signals across training differ in incorrect relative to correct trials (See Figure S15A). Green and red bars denote timepoints of significant enhancement and reduction, respectively. (**C**) Average Z-scored Ca^2+^ signals of VIP interneurons during the Delay-to-Choice transition for correct trials in NS and HFS mice in Early and Late stages of training (left). Average Ca^2+^ signals in the final 2 sec of the delay period (“distal delay”, yellow highlighted timepoints) between Early and Late training (right). Mixed-effects model, Stimulation x Stage interaction: F(1, 19)=7.135, p<0.05; *p<0.05, different from NS Early; n=9-13. NS data reproduced from A. (**D**) Functional coefficient estimates of the interaction between the Training and Stimulation covariates in correct trials, showing how changes in average Ca^2+^ signals across training differ in HFS relative to NS mice. Red bar denotes timepoints of significant reduction. (**E**) Correlation of average Z-scored VIP Ca^2+^ signals during the distal delay of correct trials in Early and Late training with overall training performance (% accuracy) in NS and HFS mice. Correlation for Early training, Overall: F(1, 19)=7.453, p<0.05; R^2^=0.2817. (**F**) Average Z-scored Ca^2+^ signals of VIP interneurons during the Delay-to-Choice transition for correct trials in NS and *Df(16)A^+/–^*mice in Early and Late stages of training (left). Average Ca^2+^ signals in the distal delay (yellow highlighted timepoints) between Early and Late training (right). Two-way ANOVA, Genotype x Stage interaction: F(1, 15)=9.657, p<0.01; *p<0.01, different from NS Early; n=8-9. NS data reproduced from C. (**G**) Functional coefficient estimates of the interaction between the Training and Genotype covariates in correct trials, showing how differences across training in the average Ca^2+^ signals differ in *Df(16)A^+/–^* relative to WT mice. Red bars denote timepoints of significant reduction. (**H**) Correlation of average Z-scored VIP Ca^2+^ signals during the distal delay of correct trials in Early and Late training with overall training performance (% accuracy) in NS and *Df(16)A^+/–^* mice. Correlation for Early training, Overall: F(1, 15)=14.73, p<0.005; R^2^=0.4955. NS data reproduced from E. (**I**) Schematic showing AAV.CaMKIIα::hChR2(H134R)-mCherry in unilateral vHPC of wildtype (WT) or *Df(16)A^+/–^* mice. (**J**) Representative images of vHPC axon terminals (yellow outlines) co-expressing mCherry (dark diffused material) and VGluT1 (gold particles, yellow arrowheads) establishing asymmetric synapses (purple arrowheads denoting postsynaptic density [PSD]) on mPFC VIP+ (left, green outline; green arrowheads indicated VIP detected by gold particles) or VIP- (right, blue outline) dendrites. (**K**) Average percent of vHPC terminals on VIP interneurons in WT and *Df(16)A^+/–^*mice (* unpaired t-test: t(10)=2.71, p<0.05; n=6,6).

### Prior vHPC stimulation selectively alters distal delay VIP interneuron activity during early learning

Having identified a training-dependent relationship between distal delay VIP interneuron activity and trial outcome, we next examined whether prior vHPC stimulation altered this activity pattern. During early training, prior HFS selectively elevated distal delay VIP interneuron activity during correct trials, abolishing the activity decrease observed in NS mice (Figure 6C). In contrast, distal delay VIP interneuron activity during incorrect trials was largely unaffected by prior HFS (Figure S15C). Consequently, VIP interneuron activity during correct trials in HFS mice resembled the activity pattern observed during incorrect trials. Consistent with these observations, training-related increases in distal delay VIP interneuron activity during correct trials were significantly blunted in HFS mice relative to NS controls (Figure 6D; S14).

Importantly, the magnitude of distal delay VIP interneuron activity during correct trials early in training negatively correlated with subsequent task performance, such that mice exhibiting higher activity showed poorer SWM acquisition (Figure 6E). In contrast, distal delay VIP interneuron activity during late training was unrelated to performance. These findings suggest that elevated VIP interneuron activity during the distal delay period in early training, promoted by prior vHPC input stimulation, is associated with less effective SWM task learning.

### Altered delay-period VIP interneuron activity is recapitulated in *Df(16)A^+/–^* mice

If elevated distal delay VIP interneuron activity contributes to impaired SWM acquisition, then a mouse model with known SWM learning deficits might exhibit a similar phenotype. To test this possibility, identical recordings were performed in *Df(16)A^+/–^* mice, a model of 22q11.2 deletion syndrome with established impairments in hippocampal-prefrontal connectivity and SWM task acquisition (Mukai et al., 2015; Passecker et al., 2026, 2025; Sigurdsson et al., 2010; Tamura et al., 2017). Paralleling their wildtype counterparts (NS mice), *Df(16)A^+/–^* mice expressing ChrimsonR in vHPC and GCaMP6f in mPFC VIP interneurons were trained on the SWM task without any prior stimulation (Figure S9).

Consistent with prior reports, *Df(16)A^+/–^* mice showed impaired SWM task acquisition relative to wildtype controls (Figure S15D). Strikingly, recapitulating the phenotype observed following prior vHPC stimulation, *Df(16)A^+/–^* mice showed elevated distal delay VIP interneuron activity during early correct trials relative to NS mice, and this activity remained elevated throughout training (Figure 6F). Consequently, *Df(16)A^+/–^* mice displayed blunted training-related increases in distal delay VIP interneuron activity in correct trials compared with NS mice (Figure 6G; Figure S15E). Furthermore, as observed in NS and HFS mice, greater distal delay VIP interneuron activity in NS and *Df(16)A^+/–^* mice during early, but not late, training predicted poorer SWM task acquisition (Figure 6H).

### *Df(16)A^+/–^* mice exhibit reduced synaptic targeting of vHPC inputs onto VIP interneurons

The convergence of behavioral and physiological phenotypes between HFS-treated and *Df(16)A^+/–^* mice raised the possibility that both manipulations alter connectivity between vHPC inputs and VIP interneurons. To test this hypothesis, electron microscopy was used to quantify the postsynaptic targets of monosynaptic vHPC inputs in wildtype and *Df(16)A^+/–^* mice (Figure 6I,J). vHPC terminals were identified in serial ultrathin sections and their postsynaptic dendrites classified as VIP-positive or VIP-negative. The proportion of vHPC synapses formed onto VIP-positive dendrites was significantly reduced in *Df(16)A^+/–^* mice relative to wildtype controls (Figure 6K). Together, these complementary findings implicate diminished connectivity between vHPC inputs and VIP interneurons as a common circuit adaptation underlying altered VIP interneuron dynamics associated with impaired SWM task acquisition.

## DISCUSSION

Here we identify a cell-type-specific form of plasticity that persistently reweights hippocampal recruitment of prefrontal inhibitory microcircuits and reshapes behaviorally relevant interneuron activity during working memory task learning. Repeated activation of vHPC inputs weakened recruitment of VIP interneurons while potentiating recruitment of SST interneurons, with more modest attenuation of PV interneuron and pyramidal neuron responses. *Ex vivo* whole-cell electrophysiology together with computational modeling identified diminished monosynaptic hippocampal drive onto VIP interneurons as a plausible circuit mechanism underlying this broader microcircuit reconfiguration. Although prior vHPC stimulation had little effect on overall SWM task acquisition, it selectively elevated distal delay VIP interneuron activity associated with successful task learning. This altered delay-period activity was independently recapitulated in *Df(16)A^+/–^* mice, which also exhibited reduced monosynaptic targeting of VIP interneurons by vHPC inputs. Together, these findings identify diminished hippocampal recruitment of VIP interneurons as a circuit mechanism through which repeated neural activity can persistently reshape prefrontal inhibitory microcircuits and their engagement during cognitive behavior.

### Activity-dependent reshaping of hippocampal recruitment across prefrontal inhibitory microcircuits

Our findings reveal robust and persistent activity-dependent plasticity in hippocampal recruitment of prefrontal inhibitory microcircuits. Rather than uniformly altering responses across neuronal populations, repeated stimulation suppressed VIP interneuron responses, enhanced SST interneuron responses, and blunted PV interneuron and pyramidal neuron responses. *Ex vivo* electrophysiology and computational modeling together support a framework in which diminished monosynaptic hippocampal recruitment of VIP interneurons propagates through canonical cortical circuitry, relieving inhibition of SST interneurons and thereby increasing inhibition of both PV interneurons and pyramidal neurons. Notably, this disinhibitory framework does not fully explain all aspects of the observed plasticity, particularly the distinct time courses and stimulation sensitivities of VIP and SST interneuron response changes (e.g., Figure 2C,G). Indeed, vHPC input stimulation likely engages multiple plasticity mechanisms beyond reduced monosynaptic drive onto VIP interneurons. Importantly, however, changes in VIP and SST interneuron responses need not scale proportionally, owing to nonlinear circuit interactions, differential amplification through recurrent networks, and differences in how population Ca²⁺ signals reflect underlying neuronal activity and synaptic transmission. Despite these complexities, relatively modest IO stimulation was sufficient to reduce hippocampal recruitment of VIP interneurons while potentiating SST interneuron responses (Figures 2D,H), consistent with the proposed disinhibitory mechanism. Collectively, these findings identify hippocampal recruitment of VIP interneurons as a particularly sensitive point of control through which activity-dependent plasticity can reconfigure prefrontal inhibitory microcircuits.

### Heightened delay-period VIP interneuron activity in early training may impair SWM task learning

By analyzing VIP interneuron dynamics from the onset of SWM task training, we identified two behaviorally relevant features of their delay-period activity: (1) activity was higher in incorrect than correct trials in early training; and (2) heightened activity during early correct trials, promoted by prior vHPC input stimulation and recapitulated in *Df(16)A^+/–^*mice, was associated with poorer SWM task acquisition. Together, these observations suggest that elevated distal delay VIP interneuron activity represents a maladaptive prefrontal microcircuit state during early learning.

One possible explanation follows from the well-established disinhibitory role of VIP interneurons within cortical microcircuits. By suppressing other inhibitory interneurons, VIP interneurons amplify pyramidal neuron activity, a mechanism thought to strengthen task-relevant cortical representations during well-learned working memory tasks (Fu et al., 2014; Kamigaki & Dan, 2017; Lee et al., 2019; Pi et al., 2013). Early in learning, however, before animals have acquired the task rule, these activity dynamics are still emerging (Meyers et al., 2012; Qi et al., 2011; Tang et al., 2022). Under these conditions, elevated VIP interneuron activity may instead amplify immature or task-irrelevant network dynamics, increasing the likelihood that these activity patterns will influence subsequent choice behavior. Conversely, lower delay-period VIP interneuron activity in early training may limit the amplification of these immature network dynamics, reducing their influence on decision-making. Consistent with this interpretation, VIP interneuron activity during the distal delay was higher in incorrect than correct trials early in training (Figure 6A,B). In our delayed non-match-to-sample task, lower delay-period VIP interneuron activity may minimize interference with decision-making, allowing mice to follow their innate tendency to explore the newly available arm (Lalonde, 2002) and thereby make the correct choice.

The same framework may also explain why elevated delay-period VIP interneuron activity specifically during early correct trials predicted poorer task learning. Early in training, correct choices often occur by chance, or through innate exploratory behavior, rather than by informed application of the task rule. Consequently, correct trials can occur before delay-period cortical activity reliably encodes task-irrelevant information. Under these conditions, elevated VIP interneuron activity may amplify immature cortical representations that happen to precede successful choices, increasing the likelihood that these representations are reinforced during learning despite being less effective at guiding future behavior. Such amplification could interfere with the normal refinement of task-relevant cortical representations across training, ultimately impairing task acquisition. In this framework, heightened delay VIP interneuron activity during early learning may maladaptively amplify task-irrelevant cortical representations, thus interfering with their normal learning-related refinement as animals acquire the task rule and ultimately impairing SWM task acquisition. These findings therefore suggest that the behavioral significance of delay-period VIP interneuron activity depends on the stage of learning and the maturity of underlying cortical representations.

### Potential neuroadaptations underlying elevated delay-period VIP interneuron activity during early learning

Unexpectedly, prior vHPC input stimulation, despite weakening monosynaptic recruitment of VIP interneurons by hippocampal input, increased distal delay-period VIP interneuron activity in early SWM training. Strikingly, *Df(16)A^+/–^* mice, whose VIP interneurons show reduced vHPC input synaptic targeting, exhibited a similar elevation in delay-period VIP interneuron activity. Together, these observations suggest that neuroadaptations downstream of diminished hippocampal recruitment of VIP interneurons underlie this behaviorally relevant activity pattern. For example, weakening vHPC input may induce compensatory reweighting of excitatory drive onto VIP interneurons by other brain regions, such as the mediodorsal thalamus (MD) (Canetta et al., 2020). MD inputs are known to directly target VIP interneurons and sustain task-relevant cortical representations across delay periods (Bolkan et al., 2017; Mukherjee et al., 2021; Schmitt et al., 2017). Thus, enhanced VIP interneuron recruitment by MD inputs could maladaptively amplify delay-period VIP interneuron activity during early training. Alternatively, neuroadaptations in prefrontal dopamine signaling may contribute to the heightened delay-period VIP interneuron activity in early learning. Dopamine has long been implicated in sustaining cortical dynamics during working memory (Arnsten, 1997; Durstewitz et al., 2000; Sawaguchi and Goldman-Rakic, 1991), and abnormally heightened mPFC D1 receptor signaling in VIP interneurons, whether arising downstream of repeated vHPC input activation (Gurden et al., 2000; Sorg et al., 2004) or through the *Df(16)A^+/–^*mutation (Choi et al., 2018), could preferentially enhance distal delay-period VIP interneuron activity (Anastasiades et al., 2018; Bae et al., 2025) and thereby impair SWM learning (Zahrt et al., 1997). The precise mechanisms of the altered delay VIP interneuron activity remain speculative and warrant future investigation.

In conclusion, we identify a form of activity-dependent plasticity that persistently reshapes hippocampal recruitment of prefrontal inhibitory microcircuits through selective weaking of monosynaptic input onto VIP interneurons. This plasticity is accompanied by altered behaviorally relevant VIP interneuron activity during working memory learning and is recapitulated in a disease-relevant model with diminished hippocampal recruitment of VIP interneurons. Together, these findings establish hippocampal recruitment of prefrontal interneurons as a dynamic and plastic circuit feature capable of influencing cortical activity during cognition and provide a framework for understanding how maladaptive reconfiguration of long-range inhibitory recruitment may contribute to cognitive dysfunction.

## MATERIALS AND METHODS

### Mice

C57BL/6J (000664), VIP::Cre (031628), SST::Cre (013044), PV::Cre (017320), SST::Flpo (031629), and Ai9-tdTomato (007909) mice were obtained from The Jackson Laboratory (Bar Harbor, ME). *Df(16)A^+/–^* mice were obtained from Dr. Joseph Gogos (Columbia University) and were backcrossed for >10 generations with C57BL/6J mice prior to transfer. *Df(16)A^+/–^* colonies were maintained by *Df(16)A^+/–^*x C57BL/6J crossings.

The following crossings were used to generate mice for different experiments: Baseline evoked response and plasticity characterization: mice homozygous for VIP, SST, or PV::Cre were crossed with C57BL/6J. Slice electrophysiology: VIP::Cre mice were crossed with SST::Flpo mice to generate VIP::Cre;SST::Flpo mice. SWM task experiments: VIP or SST::Cre mice were crossed with Ai9-tdTomato Cre reporter mice to generate Cre;Ai9-tdTomato mice; male *Df(16)A^+/–^* mice were crossed with female VIP::Cre;Ai9-tdTomato mice to generate triple-transgenic experimental mice and *Df(16)A^+/+^* controls. Electron microscopy: male *Df(16)A^+/–^* mice were crossed with female C57BL/6J mice to generate *Df(16)A^+/–^* and *Df(16)A^+/+^*controls.

Adult mice of both sexes were used (2.5-8 months at time of surgery). Mice were group-housed with littermates (up to 5 per cage) in a temperature- and humidity-controlled National Institutes of Health animal facility on a 12-h light-dark cycle. Mice were given *ad libitum* access to food and water except during food restriction for SWM training (see Spatial working memory assay). Approximately equal numbers of male and female mice were used across all experiments. All procedures were performed in accordance with the US National Research Council Guide for the Care and Use of Laboratory Animals and were approved by the National Institute of Neurological Disorders and Stroke, National Institute of Mental Health, and National Institute on Drug Abuse Animal Care and Use Committees.

### Surgical Procedures

#### Surgical Preparation

Adult mice were anesthetized with 3-5% isoflurane (v/v in oxygen, 1 L/min) in an induction chamber, weighed, shaved at the surgical site, and positioned in a stereotaxic apparatus (Kopf Instruments, Model 900) using non-rupture ear bars (Model 922). Sterile ophthalmic ointment (Paralube) was applied to prevent corneal drying. The scalp was disinfected with Betadine and 70% isopropyl alcohol. Mice were maintained at 0.8-2% isoflurane via nose cone on a 45°C heating pad (TC-1000, CWE). Depth of anesthesia was tested by toe pinch and supplemented as needed. Surgical instruments were sterilized in a hot bead sterilizer (Braintree Scientific) and cooled prior to use.

#### Virus injection

A midline scalp incision was made and the skull exposed. The skull was leveled at bregma and lambda (±0.05 µm in the D/V plane). For opto-photometry experiments: Burr holes were drilled above unilateral vHPC (in mm, A/P: −3.5, M/L: ±3.35) and ipsilateral mPFC (A/P: +1.9, M/L: ±0.4) in VIP, SST, or PV::Cre mice. AAV1.Syn::ChrimsonR-tdTomato (Addgene, 59171) or AAV1.CAG::tdTomato (Addgene, 59462), diluted in sterile saline to 5×10^12^ GC/mL, was injected into vHPC (D/V: −3.25 from brain surface; 700nL) using a pulled glass pipette connected to a Hamilton syringe (Model 75RNSYR). AAV9.Syn::FLEX.GCaMP6f (Addgene, 100833), AAV9.CaMKIIα::GCaMP6f (Addgene, 100834), or AAV5.hSyn::DIO.EGFP (Addgene, 50457) at titers of 2.5×10^12^-2.8×10^13^ GC/mL was injected into mPFC (D/V: −1.45; 500 nL). For slice electrophysiology experiments: Burr holes were drilled above bilateral vHPC (A/P: −3.5, M/L: ±3.35) and bilateral mPFC (A/P: +1.9, M/L: ±0.4) in VIP::Cre;SST::Flpo mice. AAV1.Syn::ChrimsonR-tdTomato (Addgene, 59171, 5×10^12^ GC/mL) was injected into vHPC (D/V: −3.25; 600nL). A combination of AAV9.CAG::FLEX.tdTomato (Addgene, 51503) and AAV9.EF1a::fDIO.EYFP (Vector Biolabs; final titers of 1.7×10^12^ and 1.25×10^12^ GC/mL) was injected into mPFC (D/V: −1.45, 500 nL). For SWM experiments: Burr holes were drilled above bilateral vHPC (A/P: −3.5, M/L: ±3.35) and bilateral mPFC (A/P: +1.95, M/L: ±0.7, 15° angle) in VIP::Cre;Ai9-tdTomato or SST::Cre;Ai9-tdTomato mice (with or without the *Df(16)A^+/–^* mutation). Two injections per hemisphere of AAV5.CaMKIIα::ChrimsonR-tdTomato (Vector Biolabs; 2×10^12^) were made into vHPC (D/V: −4.25 and –3.75 from bregma; 1 μL at each depth; 100 nL/min) using a syringe pump (Quintessential Stererotaxic Injector, Stoelting). AAV9.Syn::FLEX.GCaMP6f (Addgene 100833; 2.8×10^13^) was injected into mPFC (D/V: −1.8 from bregma; 500nL) ≥10 min using a Hamilton syringe and 32-gauge needle (Model 7001KH). In all cases, injectors were left in place for ≥5 min after infusion to facilitate diffusion before withdrawal. Mice used for opto-photometry and slice electrophysiology experiments had incisions sealed with tissue adhesive (Vetbond, 3M), recovered for 8-13 weeks, and then underwent optic fiber implantation. Mice used for SWM experiments underwent optic fiber implantation immediately following viral infusion.

#### Fiber implantation

For opto-photometry and slice electrophysiology experiments, holes were drilled above mPFC (A/P: +1.9, M/L: ±0.4), and two miniature screws were placed (approx. A/P: −1.0, M/L: ±2.5). After piercing dura, unilateral (opto-photometry) or bilateral (slice electrophysiology) optic fibers (200-µm core, 0.39 NA; ThorLabs) were lowered into mPFC (D/V: −1.0 from brain surface). For SWM experiments, bilateral optic fibers (200-µm core, 0.39 NA) were implanted at a 15° angle (DV: −1.6 from bregma). Fibers and screws were secured using dental cement (Unifast Trad or 3M RelyX Unicem Cement Automix; Henry Schein).

#### Postoperative procedures

Ketoprofen (5-10 mg/kg, s.c.) and sterile saline (1 mL) were administered 30 min prior to end of surgery. Mice were rehoused with one or more littermates in clean cages, except SWM mice, which were single-housed post-surgery. Ketoprofen and saline were administered 24 and 48 h post-surgery. Mice recovered for ≥10 days prior to experiments.

### Fiber photometry systems

Custom spectrometer-based systems (based on published systems (Brockway et al., 2023; Meng et al., 2018; Sciolino et al., 2022)) were used. For opto-photometry system: 473-nm blue light (MBL-III-473-100mW, Ready Lasers) passed through a Noise Eater (NEL01, ThorLabs) and 635-nm red light (MRL-III-635L-200mW, Ready Lasers), were combined using a kinematic fluorescence filter cube (DFM1, ThorLabs) onto a quadruple-edge dichroic mirror (ZT440/488/561/635rpc, Chroma). Light was coupled through an FC/PC fiber coupler (PAF2-A4A) through a fiber rotary joint (FRJ_1x1_FC-FC, Doric Lenses) and patchcords (200-µm core, 0.48 NA, Doric) to implanted fibers. Blue and red light power were calibrated to ∼70 µW and ∼5 mW output at the implanted ferrule end. Fluorescence emission was filtered (ZET488/561/633m, Chroma) and directed (PAF2S-11A, Thorlabs) into an anti-reflection-coated fiber (M200L02S-A, ThorLabs) connected to a spectrometer (QEPro, Ocean Insight). TTLs from a waveform generator (PulsePal v2, SanWorks) triggered spectral integration (17 ms) events at 40 Hz and 5-ms red laser pulses at 1-40 Hz. Spectrometer TTLs were delayed by 5 ms relative to the red laser TTLs to prevent spectral contamination. The waveform generator also delivered 20-Hz TTLs to the camera (FLIR Blackfly S USB3) to trigger the shutter. Video frames were processed using Bonsai operating real-time DeepLabCut processing nodes (Kane et al., 2020; Lopes et al., 2015) to obtain mouse body part and maze coordinates. Camera shutter events were recorded in OpenEphys for alignment of photometry and positional data. SWM photometry system: Blue light only was used with a dual-edge dichroic mirror (ZT488/561rpc, Chroma) and dual-band emission filter (ZET488/561m, Chroma). Acquisition was at 20 Hz (37-ms integration) with camera shutter TTLs synchronized to spectral integration.

### Optogenetic stimulation parameters

Opto-phometry experiments: Over 50 days, mice received six “input-output” (IO) sessions (Days 1, 8, 15, 22, 29, and 50). Each began with a 10-min baseline recording, after which vHPC terminals were stimulated with 5-ms, ∼5-mW pulses of red light across a range of pulse numbers and frequencies. IO stimulation consisted of two rounds these stimulation conditions: 5 Hz (1, 5, and 10 pulses); 10 Hz (1, 5, 10, and 20 pulses); 20 Hz (1, 5, 10, 20, and 30 pulses); and 40 Hz (1, 5, 10, 20, 30, and 40 pulses). Each stimulation condition was delivered as a 30- sec trial with 10 sec pre-stimulation and 19 sec post-stimulation. On each of 12 days early in the 50-day timeline (Days 4-7, 11-14, and 18-21), a subset of mice (“IO+HFS”) received additional “high-frequency stimulation” (HFS) of vHPC inputs consisting of 100 trains of 40 pulses at 40 Hz (30-sec inter-trial interval). “IO” mice experienced identical handling/tethering/recording but no red pulses. Mice used for the slice electrophysiology or SWM experiments received bilateral stimulation according to the 12-day HFS regimen described above or matched handling without red pulses (NS). To characterize HFS-induced response changes in mice used in the SWM experiments, on Days 1, 9 and 18, HFS mice received 50 trains of unilateral stimulation during photometry (sequentially for both hemispheres), followed by 50 trains of bilateral stimulation without photometry. NS mice received identical conditions but without red laser pulses. The hemisphere showing the most robust and characteristic GCaMP and tdTomato spectral features and the most pronounced changes in evoked Ca^2+^ response across HFS days was selected for recording during SWM task training.

### Slice electrophysiology

#### Acute brain slice preparation

Mice were anesthetized using Euthanasia solution (sodium pentobarbital and phenytoin; NIH Veterinarian Services) and decapitated. Brains were extracted into ice-cold cutting solution containing (in mM): 92 NMDG, 20 HEPES, 25 glucose, 30 NaHCO_3_, 2.5 KCl, 1.2 NaPO_4_ saturated, 10 Mg-sulfate, and 0.5 CaCl_2_ with 95% O_2_/5% CO_2_ (osmolarity 303–306 mOsm, Wescorp). The extracted brain was promptly blocked and affixed to a vibratome platform (Leica VT1200) containing ice-cold NMDG-based cutting solution. Coronal slices (250 µm) containing mPFC were made prepared (blade advance 0.07 mm/s). Slices were incubated for 5-10 min in NMDG-based solution at 34°C and then transferred to room temperature artificial cerebrospinal fluid (aCSF) saturated with 95% O_2_/5% CO_2_ for ≥1 hour before recording. aCSF contained (in mM): 92 NaCl, 20 HEPES, 25 glucose, 30 NaHCO_3_, 2.5 KCl, 1.2 NaPO_4_, 1 mM Mg-sulfate, and 2 mM CaCl_2_, with osmolarity of 303–306 mOsm.

### *Ex vivo* whole cell electrophysiology

Whole-cell patch-clamp electrophysiology studies were conducted following previously described methodology (Enriquez-Traba et al., 2024; Wang et al., 2024; Yarur et al., 2023). Cells were visualized using infrared differential interference contrast (IR-DIC) optics on an inverted microscope (Olympus BX5iWI). The recording chamber was perfused by pump (World Precision Instruments) at 1.5-2.0 mL/min with aCSF containing (in mM): 126 NaCl, 2.5 KCl, 1.4 NaH_2_PO_4_, 1.2 MgCl_2_, 2.4 CaCl_2_, 25 NaHCO_3_, and 11 glucose (303-305 mOsm). For whole-cell recordings of AMPAR- and NMDAR-mediated currents, glass microelectrodes (3–5 MΩ) were used, containing (in mM): 117 cesium methanesulfonate, 20 HEPES, 0.4 EGTA, 2.8 NaCl, 5 TEA-Cl, 4 Mg-ATP, and 0.4 Na-GTP (280-285 mOsm). SST and VIP cells were identified by tdTomato and EGFP fluorescence, respectively. Neurons were voltage-clamped using a Multiclamp 700B amplifier (Molecular Devices). Data were filtered at 2 kHz and digitized at 20 kHz with a 1440A Digidata Digitizer (Molecular Devices). Series resistance (<20 MΩ) was monitored by −5 mV voltage steps; cells with >20% change were excluded from further analysis. Monosynaptic optical EPSCs (oEPSCs) evoked by vHPC input stimulation (1-ms red light pulses of field illumination, 10-sec inter-pulse interval) were isolated using TTX (1 µM, Tocris) and 4-AP (50 µM, Tocris). For biophysically isolated AMPAR and NMDAR currents, cells were held at −70 mV and +40 mV, respectively. NMDAR current peaks were detected within 60 ms of the onset of red-light stimulation. For pharmacologically isolated AMPAR and NMDAR currents, cells were held at +40 mV and NMDAR currents were digitally subtracted from recording with the presence of D-AP5 (50 µM, Abcam).

### Spatial working memory assay

#### Maze apparatus

A custom-built, Python-controlled, automated T-maze was used. Maze arms were 12.7 cm wide and 30.5 cm high. The center arm was 60 cm long (including an 18-cm start box), and the goal arms were 26.7 cm long. The start box and goal arms contained a reward port for infrared beam-triggered delivery of sweetened condensed milk (∼40 μl, 20% in deionized water) through a blunt 20G needle. A circular rotary platform, positioned such that one half comprised the choice point of the maze, rotated 180° during delay periods to minimize use of scent cues.

#### SWM task

One day following the stimulation protocol, mice began food restriction (1.5-3g chow daily) to maintain weight at 80%-85% pre-restriction weight (adjusted for implant weight). Mice were food restricted for three days before two daily 30-min sessions of maze habituation in which all arms were milk-baited. Mice were tethered to the optical fiber on the second habituation session and all subsequent sessions. Mice underwent two or more daily sessions of shaping trials in which they visited a single available arm (randomized left/right) for milk reward, returned to the start box, and after 10 sec were forced to visit the alternate arm for reward. Shaping trials had a 20-sec inter-trial interval (ITI). Mice advanced after completing 10 trials in 30 min on two days. Mice were then trained on a delayed non-match-to-sample (DNMS) SWM task. Each trial consisted of Sample, Delay, and Choice periods. During Sample, mice visited a single arm (randomized) for reward and returned to the start box, followed by a 10-sec Delay. During Choice, both arms were available, and reward was delivered for selecting the previously unvisited arm. ITI was 20 sec. Mice performed 10 trials per session until reaching ≥70% correct performance for three consecutive days or completing 15 training days. All training occurred during the light cycle. Experimenters conducting behavior were blinded to mouse stimulation condition and *Df(16)A^+/–^* mutation.

### Photometry during optogenetic stimulation and SWM task learning

Photometry and position data were recorded throughout optogenetic stimulation sessions and SWM training. Synchronized 20- or 40-Hz pulses were delivered to the spectrometer and camera and integrated with Bonsai-based real-time DeepLabCut tracking of environmental features and mouse position, alongside OpenEphys capture of corresponding digital timestamps. Stimulation events (e.g., trial onset/offset and parameters) and maze events (e.g., choice start) were logged as network events in OpenEphys and synchronized with photometry and position data. Data acquisition was paused for 2 sec between 30-sec stimulation trials or between SWM trials (during the 20-sec ITI), bridging post- and pre-trial recordings. Pauses enabled brief opening of a Python-controlled, servo-driven clamp on the optic fiber rotary joint, allowing release of accumulated patchcord tension before reclamping. The clamp prevented light artifacts arising from rotary joint movement during recording.

### Behavioral tracking and processing

Bonsai-DLC was used for behavioral tracking (Kane et al., 2020; Lopes et al., 2015). DLC models were created by labeling 300-500 frames of videos containing the cylindrical chamber or T-maze, comprised of 10-20 frames from each of 20-30 videos of different mice. Cylinder/maze landmarks and six body parts (nose, headcap, shoulder, midpoint, hind, and base of tail) were labeled on each frame; midpoint was used for behavioral analysis. Models were trained for approximately 750,000–1,000,000 iterations, yielding confidence values of >99% in most cases. A 20-Hz camera TTL (PulsePal v2, SanWorks) triggered frame acquisition and real-time position tracking. A ≥95% confidence threshold was applied. Exported position data were down-sampled to 10 Hz, and a Kalman Filter (pykalman.github.io) was applied to estimate XY position at all timepoints, including where position values were filtered for <95% confidence. XY position was converted from pixels to cm and used to calculate speed (cm/sec).

### Photometry processing

#### Spectral processing

Spectra were inspected to confirm GCaMP6f and tdTomato expression. GCaMP and tdTomato signals were defined as 500-541 nm and 577-618 nm, respectively; photon counts within these ranges were summed to generate timeseries.

#### Opto-photometry signal processing

GCaMP6f and tdTomato photon count timeseries were linearly detrended to correct for session-long signal decay and downsampled from 40 to 10 Hz. GCaMP %ΔF/F was calculated per stimulation trial using its mean signal from the 10-sec pre-stimulation period. %ΔF/F traces were averaged across trials with identical stimulation parameters (except when assessing within-session changes). Stimulation-evoked Ca²⁺ response magnitude was defined as the peak GCaMP %ΔF/F within 5 sec of stimulation onset. tdTomato signal was low in opto-photometry experiments (given diffuse ChrimsonR-expressing vHPC inputs), precluding spectral unmixing. Motion artifacts were negligible, as mouse movement was minimal (∼1.3 cm/s during IO sessions) and control recordings (EGFP or tdTomato) showed stable signals (±2% ΔF/F; Figure S1D,E).

#### SWM photometry signal processing

In SWM experiments, VIP::Cre;Ai9-tdTomato and SST::Cre;Ai9-tdTomato mice were used to enable robust GCaMP6f and tdTomato signals and permit spectral unmixing. Raw emission spectra were linearly unmixed in R as described previously (Brockway et al., 2023; Meng et al., 2018; Sciolino et al., 2022), yielding GCaMP6f and tdTomato coefficient timeseries (Figure S6D), which were linearly detrended to correct for signal decay. To control for motion artifacts, the ratio of unmixed GCaMP6f to tdTomato coefficients was calculated (Figure S6D). Ratio timeseries were downsampled from 20 to 10 Hz, aligned to DLC position and OpenEphys maze events, and segmented into trials. Z-scores (ZS) were computed per trial using each trial’s mean and standard deviation, yielding the Ca²⁺ signal. Ca²⁺ signals were aligned to discrete task events (sample start/end, delay, choice start/end) and averaged across trials. VIP interneuron Ca²⁺ activity was quantified as the mean ZS in the 2-sec window preceding choice start (distal delay). To assess learning-related changes, trials were divided into early (first third) and late (final third) phases for each mouse. For visual comparison of task-related VIP and SST interneuron Ca²⁺ signals and mouse speed (Figure S6E), ZS Ca²⁺ signals and speed (cm/s) were normalized to a ±1 range within individual trial events and averaged per mouse. Cross-correlations between Ca²⁺ signals and speed were computed on full trials using a maximum lead/lag of ±50 samples (±5 sec at 10 Hz; Figure S6F).

### Electron Microscopy

#### Tissue processing and immunolabeling

*Df(16)A^+/–^* and littermate wildtype mice were injected with AAV5.CaMKIIα::hChR2(H134R)-mCherry into the right vHPC. Vibratome sections (40 μm) were rinsed with phosphate-buffered saline (PBS), incubated with 1% sodium borohydride in PBS for 30 min, rinsed in PBS, and incubated with blocking solution (1% normal goat serum [NGS], 4% BSA in PBS supplemented with 0.02% saponin) for 30 min. Sections were incubated for 24 h at 4°C with primary antibodies: mouse anti-mCherry antibody (1:1000), guinea pig anti-VGluT1 antibody (1:500), and rabbit anti-VIP antibody (1:1000, 20077, ImmunoStar), all diluted in blocking solution. Sections were rinsed and incubated overnight at 4°C in the corresponding secondary antibodies: biotinylated anti-mouse (mCherry detection), anti-guinea pig-IgG Fab′ fragment coupled to 1.4-nm gold (VGluT1 detection, 1:100, 2055-1ML, Nanoprobes Inc.), and anti-rabbit-IgG coupled to 1.4-nm gold (VIP detection, 1:100, 2004-1ML, Nanoprobes Inc.). Sections were rinsed in PBS and incubated in avidin-biotinylated horseradish peroxidase complex in PBS for 2 h at room temperature. Sections were rinsed in PBS and postfixed with 1.5% glutaraldehyde at room temperature for 10 min. Sections were rinsed again in PBS and then in double-distilled water, followed by silver enhancement of gold particles with the Nanoprobe Silver Kit (2012, Nanoprobes Inc.) for 7 min at room temperature. Peroxidase activity was detected with 0.025% 3,3’-diaminobenzidine (DAB) and 0.003% H_2_O_2_ in PBS for 5-10 min. Sections were rinsed with PBS and fixed with 0.5% osmium tetroxide in PBS for 25 min, washed in PBS followed by double distilled water, and contrasted in freshly prepared 1% uranyl acetate for 35 min. Sections were dehydrated through a series of graded alcohols and propylene oxide, and flat embedded in Durcupan ACM epoxy resin (14040, Electron Microscopy Sciences). Resin-embedded sections were polymerized at 60°C for 2 days, and 60-nm sections were cut from the outer surface of the tissue with an ultramicrotome UC7 (Leica Microsystems) using a diamond knife (Diatome). Sections were collected on formvar-coated single slot grids and counterstained with Reynolds lead citrate to be examined and photographed using a Tecnai G2 12 transmission electron microscope (Fei Company) equipped with the OneView digital micrograph camera (Gatan).

#### Ultrastructural analysis

Serial ultrathin mPFC sections were analyzed blind to genotype. Synaptic contacts were classified according to morphology and immunolabel and imaged at 6,800-13,000x. Morphological criteria used for identification and classification of cellular components or type of synapse were as previously described (Peters et al., 1991). A terminal containing ≥3 immunogold particles was considered immuno-positive. Images were adjusted to match contrast and brightness using Adobe Photoshop (Adobe Systems). Experiments were repeated three times.

### Histology

Following experiments, mice were anesthetized with isoflurane (5% in oxygen, v/v) and transcardially perfused with 10 mL PBS followed by 10 mL of 4% paraformaldehyde (PFA, FD Neurotechnologies). Brains were post-fixed for 24-48 h, transferred to PBS, and sectioned at 50 μm on a vibratome (Camden Instruments 7000). Slices were mounted with DAPI Fluoromount-G (Southern Biotech), and imaged on epifluorescence (Leica), confocal (LSM800, Zeiss), or slide-scanning (Axioscan 7, Zeiss) microscopes to verify fiber placements and virus expression. Placements were mapped to atlas templates (Franklin & Paxinos, 2007).

### Experimental design and sample sizes

Sample sizes (n) indicate number of mice unless otherwise stated. Exact n for each analysis are reported in corresponding figure legends. Baseline evoked response characterization: VIP::Cre, n=23; SST::Cre, n=25; PV::Cre, n=9. Plasticity characterization: VIP::Cre, n=23; SST::Cre, n=25; PV::Cre, n=16; C57BL/6J (CaMKIIα), n=10. Overall IO/IO+HFS, n=30/44. Slice electrophysiology: VIP::Cre;SST::Flp NS/HFS, n=13/11. SWM: VIP::Cre;Ai9 NS/HFS, n=12/13; SST::Cre;Ai9 NS/HFS, n=12/15; VIP::Cre;Ai9;*Df(16)A^+/–^*NS, n=9. Electron microscopy: WT, n=6; *Df(16*)*A^+/–^*, n=6. Mice were excluded from analysis if premature death, adverse health effects, or loss of an implanted ferrule occurred prior to study completion.

### Statistical analyses

Analyses were performed using GraphPad Prism and custom Python/R scripts. For opto-photometry and SWM experiments, 2- or 3-way repeated measures ANOVA tested between-subject factors (e.g., stimulation condition) and within-subject factors (e.g., day). Mixed-effects models were conducted in Prism in rare instances with minor and random datapoint loss. Post hoc significance (p<0.05) was determined by Tukey’s/Dunnet’s multiple comparisons tests. Pairwise comparisons (e.g., NS vs. HFS oEPSCs) used Mann-Whitney or unpaired t tests depending on sample size and distribution normality. Kolmogorov-Smirnov tests compared cumulative distributions of Days to Criterion. Pearson correlations quantified relationships between Ca^2+^ signals and behavior. Within-session changes in evoked Ca^2+^ responses were tested using linear mixed-effects models (“lmer” in R; fixed effect trial; random intercept and slope per mouse). Unless stated otherwise, error bars and shaded bands represent standard error of the mean.

### Functional linear mixed modeling

Functional linear mixed modeling (FLMM) was performed using the fastFMM toolkit (Cui et al., 2022; Loewinger et al., 2024), implemented in R.

First, FLMM was used to quantify differences in IO stimulation-evoked Ca^2+^ responses in VIP, SST, and PV interneurons and putative pyramidal neurons in HFS mice on Days 1 and 8 (Figure 2A). Pairs of 40-pulse/40-Hz trials from each of Day 1 and 8 were averaged within mouse and modeled as a function of the fixed effect of Day with a random intercept for each mouse (photometry ∼ day + (1 | mouse_id)). Specifically, for animal *i*, we modeled the photometry response at time *t* as:

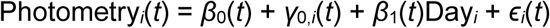

where *ɛ_i_*(*t*) ∼ N(0, *σ*^2^(*t*)) and *γ*_0,*i*_(*t*) ∼ N(0, *σ*^2^ (*t*)). We denoted Day*_i_* = 0 for Day 1 Ca^2+^ responses and Day*_i_* = 1 for Day 8 Ca^2+^ responses. This model was fit to a dataset of VIP, SST, PV, or pyramidal neuron Ca^2+^ responses (Figure 2D,G,J,M).

FLMM was also used to quantify differences in IO stimulation-evoked Ca^2+^ responses in VIP, SST, PV interneurons in IO and IO+HFS mice on Days 1 and 8 (Figure 2A). Pairs of 40-pulse/40-Hz trials from each of Day 1 and 8 were averaged within mouse and modeled as a function of two fixed effects (Day and Stimulation) and their interaction with a random intercept for each mouse (photometry ∼ day * stim + (1 | mouse_id)). Specifically, for animal *i*, we modeled the photometry response at time *t* as:

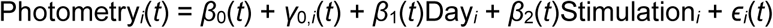

where *ɛ_i_*(*t*) ∼ N(0, *σ*^2^(*t*)) and *γ*_0,*i*_(*t*) ∼ N(0, *σ*^2^ (*t*)). We denoted Day*_i_* = 0 for Day 1 Ca^2+^ responses and Day*_i_* = 1 for Day 8 Ca^2+^ responses. We denoted Stimulation*_i_* = 0 for IO mice and Stimulation*_i_* = 1 for IO+HFS mice. This model was fit to a dataset of VIP, SST, or PV interneuron Ca^2+^ responses (Figure S4B,D,F).

Individual SWM task event-level Ca^2+^ signals spanning 10-sec windows were analyzed using five FLMMs:

The first (1) modeled Ca^2+^ signals as a function of the fixed effect of Stimulation with a random intercept for each mouse (photometry ∼ stim + (1 | mouse_id)). Specifically, for animal *i*, on event *e*, we modeled the Ca^2+^ signals at time *t* as:

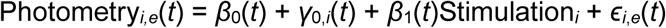

where *ɛ_i_*_,*e*_(*t*) ∼ N(0, *σ*^2^(*t*)) and *γ*_0,*i*_(*t*) ∼ N(0, *σ*^2^*_γ_*(*t*)). We denoted Stimulation*_i_* = 0 for NS mice and Stimulation*_i_* = 1 for HFS mice. This model was fit to datasets of VIP or SST interneuron Ca^2+^ signals from either correct trials (Figure 7B) or incorrect trials (data not shown).

The second (2) modeled Ca^2+^ signals as a function of the fixed effect of Outcome with a random slope and intercept for each mouse (photometry ∼ outcome + (outcome | mouse_id)). Specifically, for animal *i*, on event *e*, we modeled the Ca^2+^ signals at time *t* as:

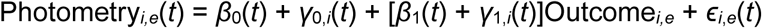

where *ɛ_i,e_*(*t*) ∼ N(0, *σ*^2^(*t*)) and *γ_i,e_*(*t*) ∼ N(0,Σ*_γ_*(*t*)). We denoted Outcome*_i,e_* = 0 for Correct trials and Outcome*_i,e_* = 1 for Incorrect trials. This model was fit to datasets of VIP or SST interneuron Ca^2+^ signals from either NS or HFS mice (Figure S6E).

The third (3) modeled Ca^2+^ signals as a function of two fixed effects (Training and Outcome) and their interaction with a random slope and intercept for Outcome for each mouse (photometry ∼ training * outcome + (outcome | mouse_id)). Specifically, for animal *i*, on event *e*, we modeled the Ca^2+^ signals at time *t* as:

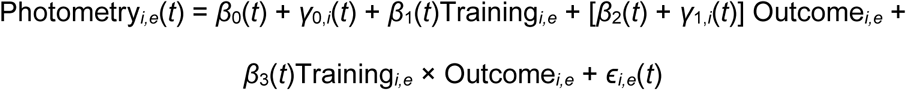

where *ɛ_i,e_*(*t*) ∼ N(0, *σ*^2^(*t*)) and *γ_i,e_*(*t*) ∼ N(0,Σ*_γ_*(*t*)). We analyzed Training*_i,e_* as a continuous variable, where 0 corresponds to the first trial and 1 corresponds to the last trial of a given mouse. We denoted the binary variable Outcome*_i,e_* = 0 for Correct trials and Outcome*_i,e_* = 1 for Incorrect trials. This model was fit to a dataset of VIP interneuron Ca^2+^ signals from NS mice (Figure 8B, S7A).

The fourth (4) modeled Ca^2+^ signals (or speed) as a function of two fixed effects (Training and Stimulation) and their interaction with a random intercept for each mouse (photometry ∼ training * stim + (1 | mouse_id)). Specifically, for animal *i*, on event *e*, we modeled the Ca^2+^ signals (or speed) at time *t* as:

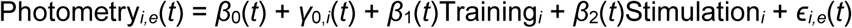

where *ɛ_i,e_*(*t*) ∼ N(0, *σ*^2^(*t*)) and *γ_i,e_*(*t*) ∼ N(0,Σ*_γ_*(*t*)). We analyzed Training*_i_* as a continuous variable, where 0 corresponds to the first trial and 1 corresponds to the last trial of a given mouse. We denoted Stimulation*_i_* = 0 for WT mice and Stimulation*_i_* = 1 for HFS mice. This model was fit to a dataset of VIP interneuron Ca^2+^ signals (or speed) from VIP::Cre NS and HFS mice (Figure 8D, S7A, S8, S9).

The fifth (5) model was identical to the fourth, except that we replaced Stimulation*_i_* with Genotype*_i_* (photometry ∼ training * genotype + (1 | mouse_id)), and denoted Genotype*_i_* = 0 for WT (NS) mice and Genotype*_i_* = 1 for *Df(16)A^+/–^* mice. This model was fit to a dataset of VIP interneuron Ca^2+^ signals from NS and *Df(16)A^+/–^*mice (Figure 8G, S7D).

Lastly, FLMM was used to quantify differences in the Ca^2+^-speed cross-correlations of SST and VIP interneuron recordings from WT, NS mice. Individual trial-level cross-correlations spanning ±5 sec lead/lag timepoints were modeled as a function of the fixed effect of CellType with a random intercept for each mouse (cc ∼ cell_type + (1 | mouse_id)). Specifically, for animal *i*, on trial *j*, we modeled the cross-correlations at time *t* as:

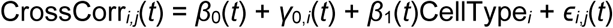

where *ɛ_i_*_,*j*_(*t*) ∼ N(0, *σ*^2^(*t*)) and *γ*_0,*i*_(*t*) ∼ N(0, *σ*^2^ (*t*)). We denoted CellType*_i_* = 0 for SST-speed cross-correlations and CellType*_i_* = 1 for VIP-speed cross-correlations. This model was fit to a dataset of Ca^2+^-speed cross-correlations from NS mice (Figure S6G).

Prior to selection, models were tested using different random effects structures; models that successfully converged and yielded lower AIC/BIC values were selected. FLMM generates Pointwise and Joint 95% confidence intervals (CIs) around the resulting functional coefficients. Timepoints at which Pointwise 95% CIs do not contain Y = 0 indicate individual timepoints of statistical significance, without correction for multiple comparisons. Joint 95% CIs, generated by exploiting the correlation between neighboring timepoints, are adjusted for multiple comparisons; thus, time intervals during which Joint 95% CIs do not contain 0 are interpreted as statistically significant.

### Computational Modeling

An integrate-and-fire computational model simulated a mPFC circuit composed of VIP, SST, PV, and pyramidal (PYR) neurons responding to vHPC input stimulation. The model included four VIP cells, six SST cells, six PV cells, and twenty PYR cells with connection probabilities and synaptic weights informed by experimental data (e.g., Figures 4E,F) and literature (Canetta et al., 2020; Marek et al., 2018b; Pfeffer et al., 2013; Sánchez-Bellot et al., 2022). Each cell type received inputs from the vHPC, mPFC PYR cells, and a subset of mPFC interneuron types. Synaptic weights also reflected the canonical view of “disinhibitory” VIP inhibiting other interneurons (predominantly SST interneurons), SST cells as providing “feedback” inhibition, and PV cells as providing “feedforward” inhibition (Fu et al., 2014; Lee et al., 2013; Pi et al., 2013).

The synaptic weights used in the model were as follows:

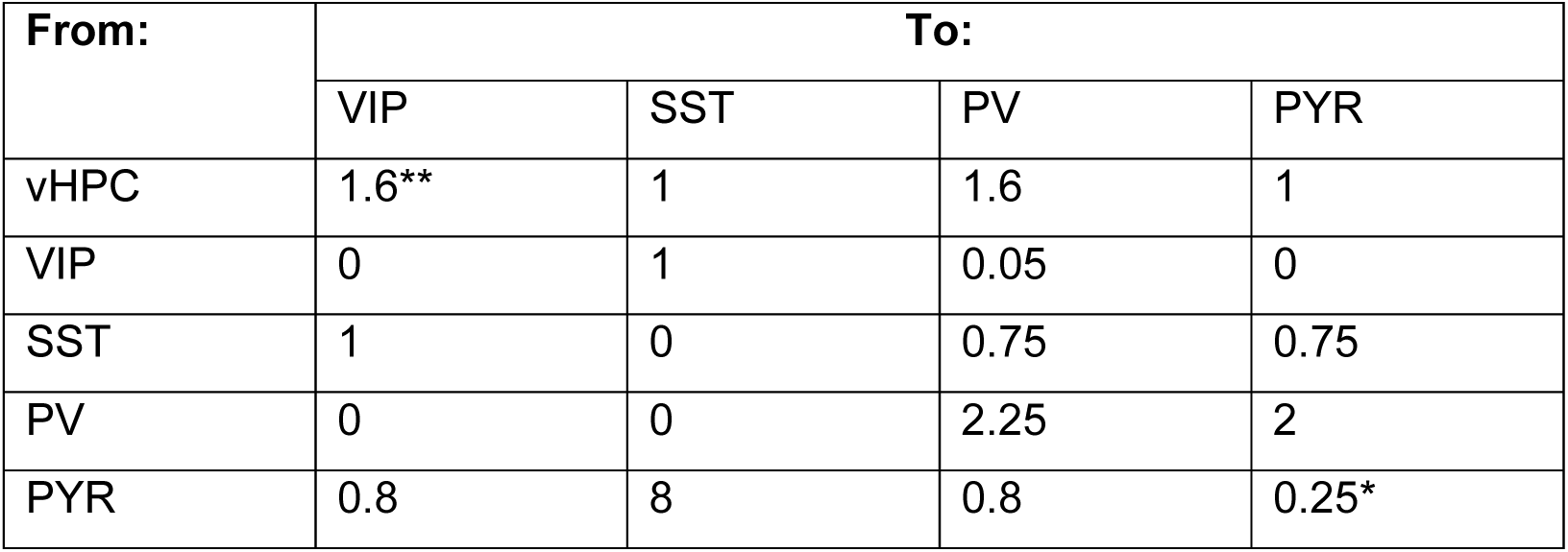

| From: | To: |  |  |  |
| --- | --- | --- | --- | --- |
|  | VIP | SST | PV | PYR |
| vHPC | 1.6** | 1 | 1.6 | 1 |
| VIP | 0 | 1 | 0.05 | 0 |
| SST | 1 | 0 | 0.75 | 0.75 |
| PV | 0 | 0 | 2.25 | 2 |
| PYR | 0.8 | 8 | 0.8 | 0.25* |

\*Not all PYR cells synapsed onto all other PYR cells; connection probability between any two PYR cells was 0.2.
\*\*Value was later modified to explore whether decreased vHPC inputs to VIP cells might plausibly reproduce dynamics we observed experimentally (e.g., Figure 2).

Network activity was simulated over the course of a 10-sec period, with vHPC input stimulation occurring from time = 2 to 3 sec. Each neuron in each cell class began with a membrane voltage of 0, and at every subsequent timestep (dt = 0.1 ms) the new membrane voltage of each neuron was calculated based on the current activity levels of each of its inputs, multiplied by the corresponding synaptic weight. Specifically, the following equation was used to estimate the membrane potential of neuron *i* from cell class *x* at timepoint *t*:

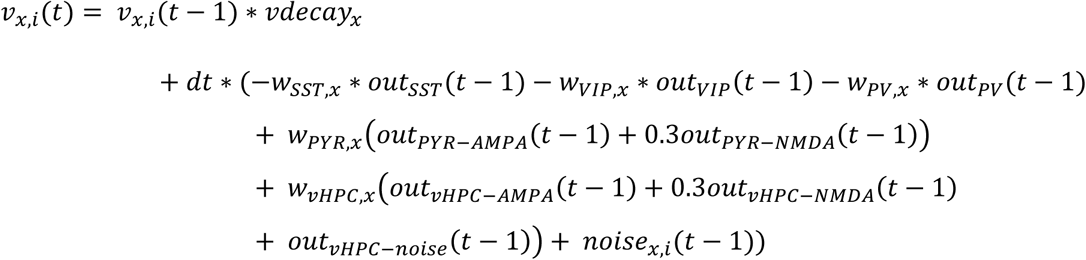

where *w_A_*_,*x*_ is the synaptic weight of neurons from cell class *A* onto neurons in cell class *x*; *out_A_*(*t*) represents the synaptic output of cell class *A* at time *t*; *noise_x_*_,*i*_(*t*) is the random noise input which differs between individual neurons from cell class *x*; and *vdecay_x_* _= *e*−*dt*/*τ*_*_x_* describes the decay of a neuron’s membrane potential back to its resting potential over time, with a time constant τ = 10 ms for all cell types. The amplitudes of NMDA outputs were scaled by a factor of 0.3, as informed by the slice electrophysiology data (Figure 4F,H). Whenever a neuron’s membrane potential crossed a threshold of 10, it was said to have fired a spike, and the membrane potential was reset to −10 at the next timestep.

For PYR neurons, this equation was modified to:

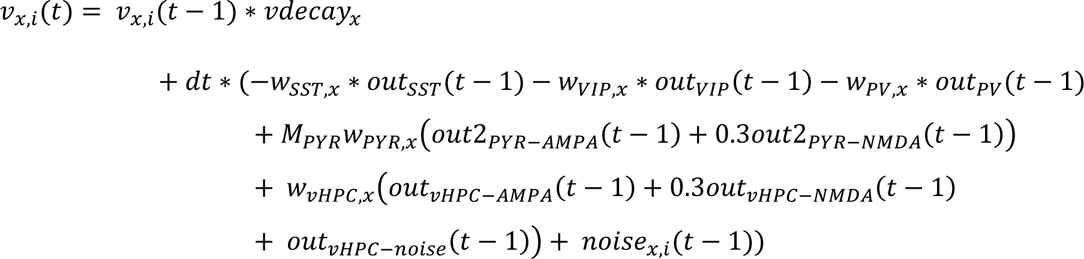

where *M_P_*_Y*R*_is a 20×20 connectivity matrix between the twenty PYR cells (connection probability between any two cells = 0.2), and *out2* is a 20×1 matrix describing the synaptic output of individual PYR cells.

To determine the synaptic output from a given mPFC cell class (*out_PV_*, *out_SST_*, *out_VIP_*, *out_P_*_Y*R*−*AMPA*_, or *out_P_*_Y*R*−*NMDA*_) at a given timepoint *t*, the following steps were performed. At t = 0, all mPFC synaptic outputs were set to 0. At each subsequent timestep, the new membrane voltages were calculated using the equation above. Next, the proportion of cells from each cell class whose membrane potential had crossed the firing threshold (threshold = 10) was determined; this proportion was called *P_x_*. The synaptic output of a cell class *x* was then calculated as:

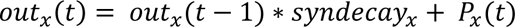

where the time constants τ for synaptic decay were 20 ms for all interneuron classes, 10 ms for PYR AMPA currents, and 80 ms for PYR NMDA currents. The same equation was used to calculate each element of the 20×1 *out2* matrix, with P_x_ simply being 1 if that PYR cell had fired and 0 otherwise.

To determine the vHPC input synaptic output at a given timepoint *t*, a raster of vHPC input spike events was created. From time = 0 to 2 sec, the rate of vHPC input firing was 0 Hz. From time = 2.0001 to 3 sec (i.e., during vHPC input ‘stimulation’), the expected rate of vHPC input firing was 100 Hz, meaning that the expected number of events within 1 sec was 100 (although the exact timing of the events was random). After time = 3 sec, the vHPC input event rate gradually decayed down from 100 Hz, with a time constant τ of 1000 ms. To determine the AMPA component of the vHPC input synaptic output *out_vHPC_*_−*AMPA*_(*t*), the event raster was smoothed by allowing each event (of amplitude 1) to decay over 40 timesteps with a time constant τ of 8 ms; the value of the smoothed trace at any timepoint *t* could then be found. To determine the NMDA component of the vHPC input synaptic output *out_vHPC_*_−*NMDA*_(*t*), the event raster was smoothed by allowing each event to decay over 400 timesteps using a time constant τ of 80 ms. Finally, all cell types also received additional noise input from the vHPC input, *out_vHPC_*_−*noise*_(*t*), representing background vHPC input that occurs even in the absence of vHPC input stimulation. Noise events occurred at an expected rate of 100 Hz throughout the entire 10-sec duration of the simulation, had an event amplitude of 1, and decayed with a time constant τ of 8 ms over 40 timesteps.

Each neuron also received neuron-specific noise, *noise_x_*_,*i*_(*t*). Random noise events for each neuron occurred at an expected rate of 100 events/s, had an amplitude of 1 (except for PV cells, where the amplitude was scaled to 1.2), and decayed over 40 timesteps with a time constant τ of 8 ms.

During each run of the model, the total number of spikes from each cell class at each timepoint was recorded. After the run, these traces of summed spike counts were converted to photometry-like activity traces by smoothing with a Gaussian filter with sigma = 25 msec. The final activity traces from each simulation were the average of five runs.

To test the hypothesis that decreased vHPC input onto VIP interneurons might explain the observed dynamics (Figure 2), the value of *w_vHPC_*_,*VIP*_ was systematically decreased from its baseline value of 1.6. Ten simulations were run for each synaptic weight value, with each simulation consisting of the average of five runs of the model. Means and SEMs were calculated using the output from the ten simulations (n = 10). To look at vHPC input stimulation-‘evoked’ activity, baseline activity for each cell type was calculated as the mean of the activity traces from time = 0.1 to 2 sec and was subtracted from the overall activity trace.

## ACKNOWLEDGEMENTS

We thank all members of the Gordon laboratory for their generous discussions, feedback, and technical advice. In particular, we thank Dr. Maxym Myroshnychenko for foundational contributions to lab infrastructure, Dr. Ako Ikegami for analysis expertise, Dr. Michelle Frazer for support with the computational model, Chloe Aloimonos for contributing to the establishment of behavioral tracking, Kirsten Gilchrist for assistance with pilot photometry experiments, and Marjorie Sapphire Bowen-Kauth for assistance with histology. We thank Dr. Guohong Cui for advice on photometry system design and code for spectral unmixing. We thank George Dold and colleagues in the NIMH Section on Instrumentation for their expert design and support of the T-maze apparatus. We thank Dr. Vincent Schram and colleagues in the NICHD Imaging Core for their training and assistance in confocal imaging. We thank the Porter Neuroscience Research Center Animal Care Staff for their exceptional animal husbandry. We thank the Genetically Encoded Neuronal Indicator and Effector (GENIE) Project and the Janelia Farm Research Campus of the Howard Hughes medical Institute for the GCaMP6f construct. This work was supported by the Division of Intramural Research of the NINDS (ZIA NS003168), NIMH (ZIA MH002970, ZIC-MH002968), and NIDA (ZIA DA000511).

## RESOURCE AVAILABILITY

Further information and requests for resources, reagents, data, and code should be directed to and will be fulfilled by David Kupferschmidt.

## AUTHOR CONTRIBUTIONS

Conceptualization, D.A.K., J.A.G., S.E.S., R.M.M., T.T.C.

Methodology, D.A.K., S.E.S., T.T.C., R.M.M., V.S.S., S.N., G.L.

Investigation and data analysis, D.A.K., S.E.S., T.T.C., M.S.D, E.V., S.N., H.E.Y., V.S.T., S.Z., R.Y., R.M.M., M.H., A.B.

Funding acquisition, F.P., M.M., V.S.S., H.A.T., J.A.G. Project administration, D.A.K., S.E.S.

Supervision, D.A.K., F.P., M.M., H.A.T., J.A.G.

Writing – original draft, S.E.S., D.A.K.

Writing – review & editing, S.E.S., D.A.K., J.A.G., V.S.S., H.A.T., R.M.M., T.T.C.

The contributions of the NIH authors were made as part of their official duties as NIH federal employees, are compliant with agency policy requirements, and are considered Works of the United States Government. However, the findings and conclusions presented in this paper are those of the author(s) and do not necessarily reflect the views of the NIH or the U.S. Department of Health and Human Services.

