## Supplemental Figures for "Activity-induced weakening of hippocampal input to prefrontal VIP interneurons reshapes inhibitory microcircuit function: implications for spatial working memory task acquisition"

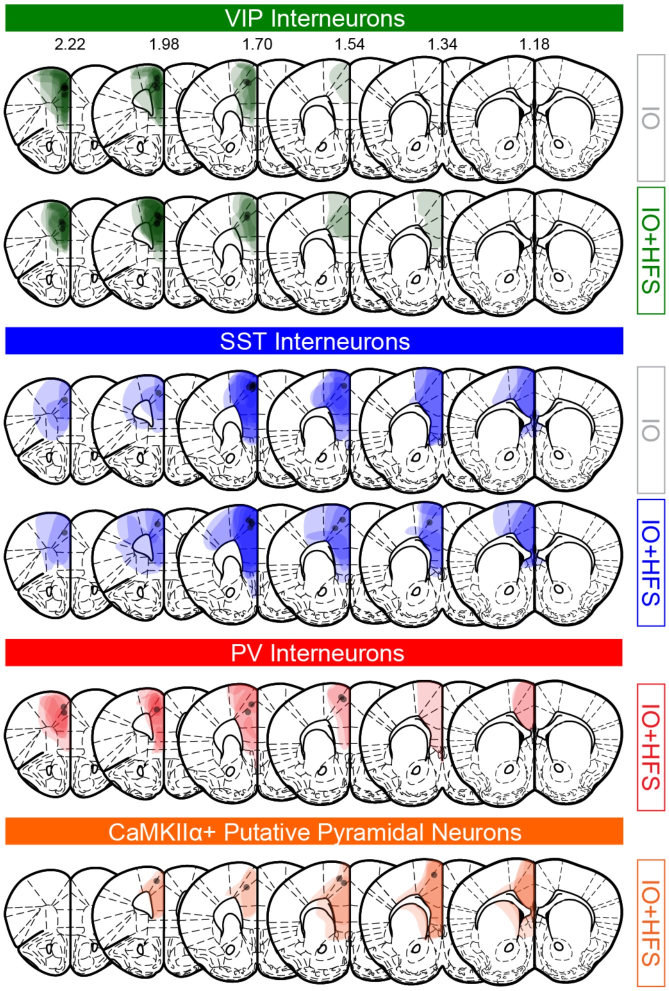


**Figure S1. Histology of GCaMP6f expression and optrode placement in mPFC for opto-photometry experiments.**

AAV.Syn::FLEX.GCaMP6f or AAV.CaMKIIα::GCaMP6f expression in mPFC VIP, SST, or PV interneurons or CaMKIIα+ putative pyramidal neurons in IO and IO+HFS mice. Gray circles indicate optical fiber placement.

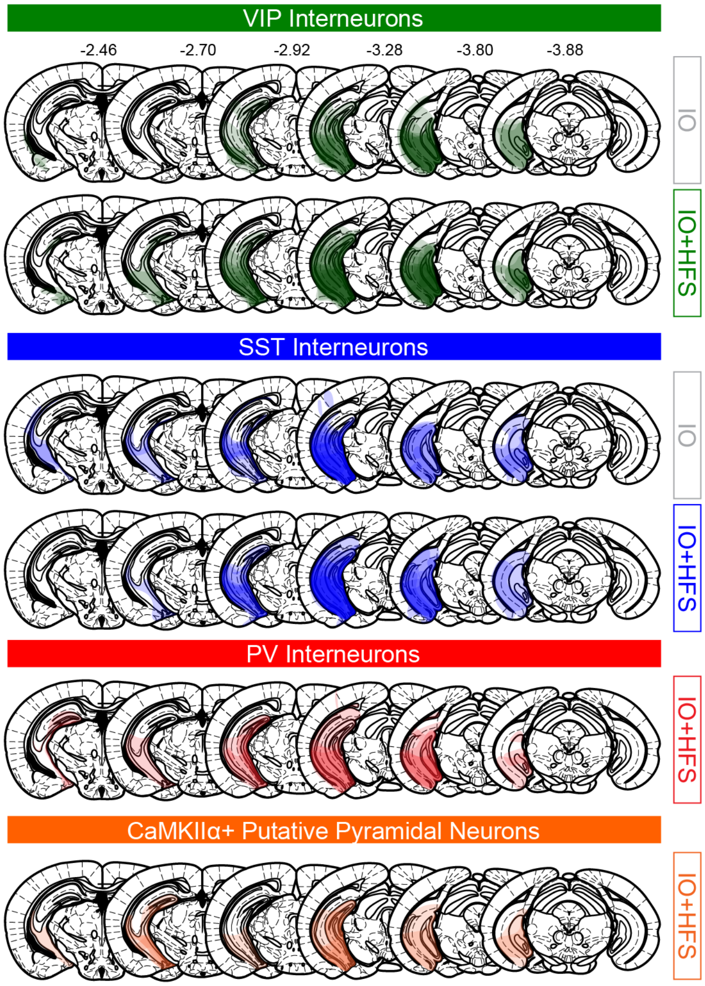


**Figure S2. Histology of ChrimsonR-tdTomato expression in vHPC for opto-photometry experiments.**

AAV.Syn::ChrimsonR-tdTomato expression in vHPC in IO and IO+HFS mice used for opto-photometry recordings from VIP, SST, or PV interneurons or CaMKIIα+ putative pyramidal neurons.

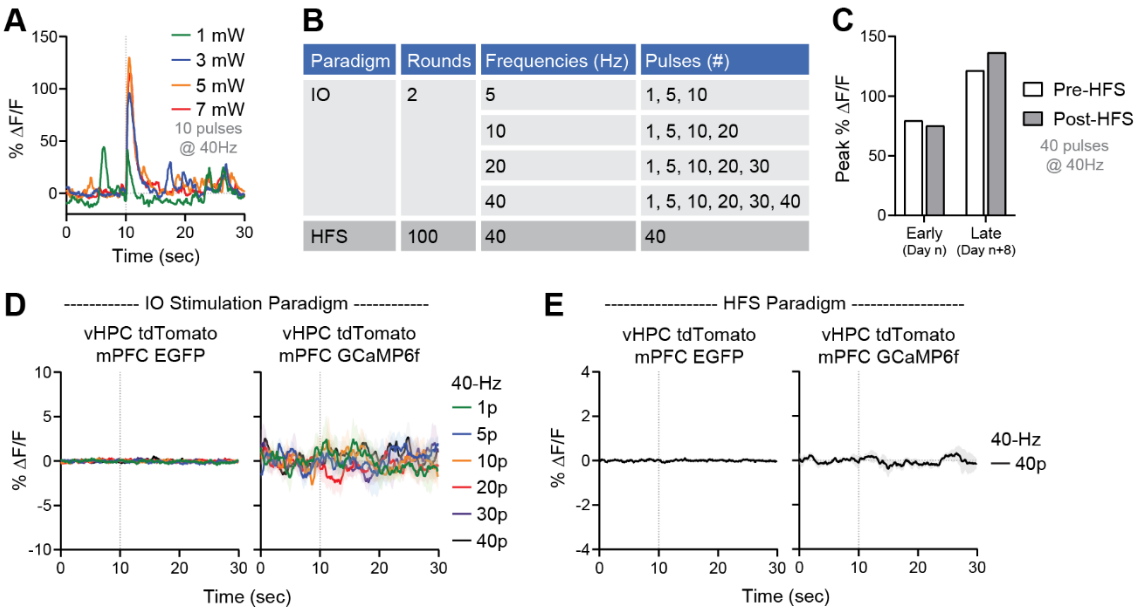


**Figure S3. Optimization and validation of the opto-photometry paradigm in SST::Cre mice.**

(**A**) Stimulation-evoked $Ca^{2+}$ responses from a representative pilot SST::Cre mouse to different red laser outputs (i.e., different vHPC input stimulation intensities). The SST::Cre mouse had previously received vHPC input stimulation and thus showed strong $Ca^{2+}$ responses. (**B**) Table description of rounds, frequencies (Hz), and pulse numbers of redlight stimulation delivered during IO and HFS sessions. (**C**) Peak stimulation-evoked $Ca^{2+}$ responses from a representative pilot SST::Cre mouse with prior vHPC input stimulation at different stages of a pilot experiment. In this experiment, peak responses to vHPC input stimulation were assessed before and after 100 trains of HFS each day for 9 days. (**D**) Average SST interneuron photometry recordings during IO stimulation (various pulse numbers at 40 Hz) in mice expressing control fluorophores. Left: Mice (n=4) expressing tdTomato in vHPC and EGFP in mPFC SST interneurons. Right: Mice (n=4) expressing tdTomato in vHPC and GCaMP6f in mPFC SST-INs. (**E**) Average SST interneuron photometry recordings during HFS stimulation in mice expressing control fluorophores. Left: Mice (n=4) expressing tdTomato in vHPC and EGFP in mPFC SST interneurons. Right: Mice (n=4) expressing tdTomato in vHPC and GCaMP6f in mPFC SST interneurons.

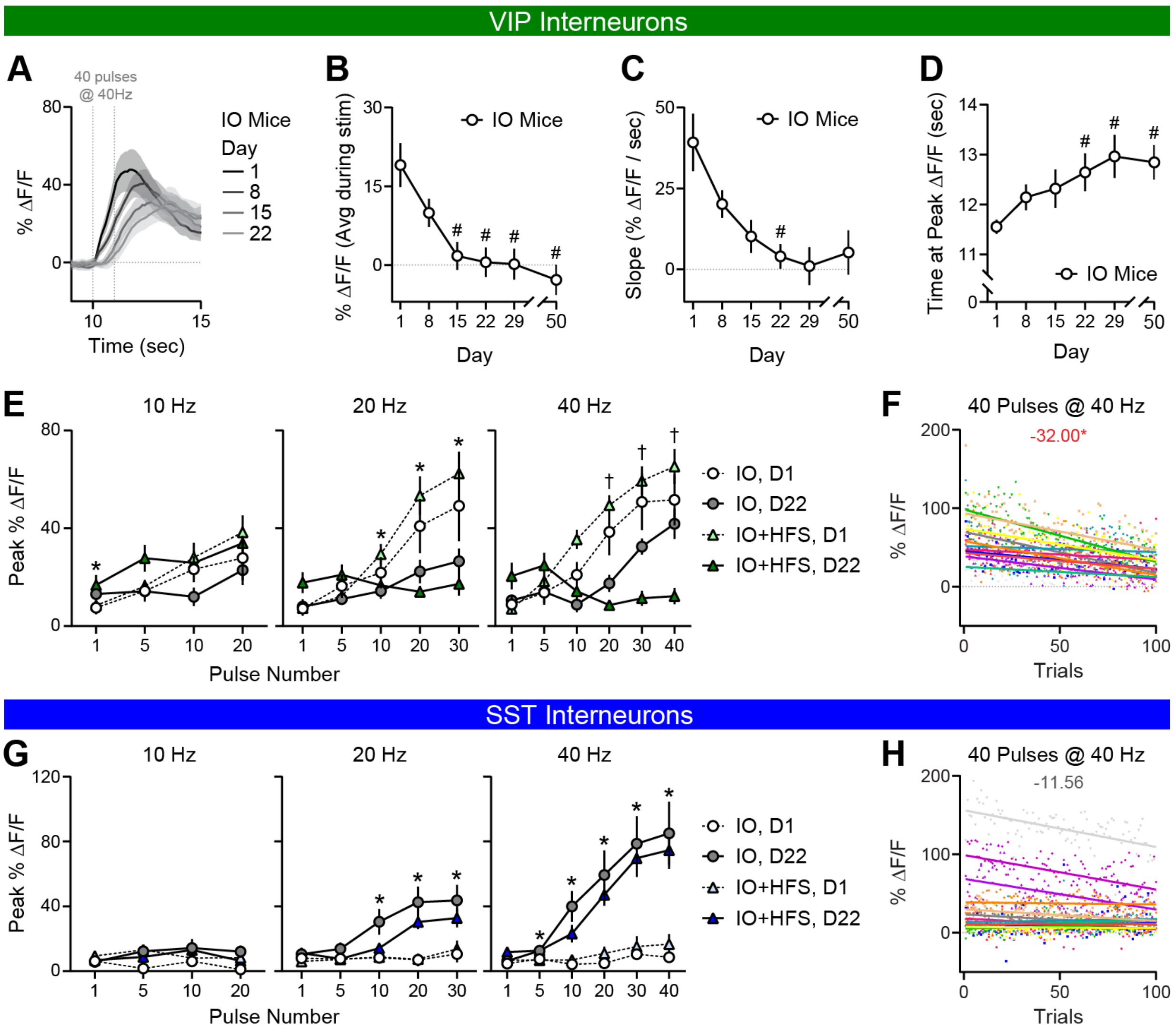


**Figure S4. Longitudinal input-output relationships and within-session response stability in VIP and SST interneurons following vHPC input stimulation.**

(**A**) Average stimulation-evoked $Ca^{2+}$ responses of mPFC VIP interneurons on Days 1, 8, 15, and 22 in IO mice (n=11). Stimulation (40 pulses at 40 Hz) initiated at 10-sec timepoint. (**B**) Average stimulation-evoked $Ca^{2+}$ responses during 1-sec stimulation (40 pulses at 40 Hz) on Days 1, 8, 15, 22, 29, and 50. Repeated measures ANOVA, Main effect of Day: F(2.309, 23.09)=8.394, p<0.005; #p<0.05, different from Day 1; n=11. (**C**) Slope of average stimulation-evoked $Ca^{2+}$ responses across 1-sec stimulation on Days 1, 8, 15, 22, 29, and 50. Repeated measures ANOVA, Main effect of Day: F(1.455, 14.55)=6.582, p<0.05; #p<0.05, different from Day 1; n=11. (**D**) Time at peak $Ca^{2+}$ response to stimulation on Days 1, 8, 15, 22, 29, and 50. Repeated measures ANOVA, Main effect of Day: F(3.321, 33.21)=4.145, p<0.05; #p<0.05, different from Day 1; n=11. (**E**) Average peak stimulation-evoked $Ca^{2+}$ activity in VIP interneurons at varying frequencies and pulse numbers in IO and IO+HFS mice on IO Days 1 and 22. 3-way ANOVA, 10 Hz: Pulse Number x Day interaction: F(2.735, 57.43)=4.000, p<0.05;

*$p<0.05$, Day 1 vs. Day 22. 20 Hz: Pulse Number x Day interaction: $F(1.364, 28.64)=20.63$, $p<0.0001$; Pulse Number x Stimulation x Day interaction: $F(4, 84)=4.622$, $p<0.005$; *$p<0.05$, Day 1 vs. Day 22. 40 Hz: Pulse Number x Stimulation interaction: $F(5, 105)=3.529$, $p<0.01$; Pulse Number x Day interaction: $F(2.592, 54.42)=19.51$, $p<0.0001$; Stimulation x Day interaction: $F(1, 21)=4.613$, $p<0.05$; Pulse Number x Stimulation x Day interaction: $F(5, 105)=6.475$, $p<0.0001$; †$p<0.005$, HFS Day 1 vs. HFS Day 22; n=11-12. (**F**) Peak stimulation-evoked $Ca^{2+}$ activity in VIP interneurons for all 100 trials on HFS Day 4. Linear Mixed Models; Fixed effect of Trial reported in red, *: $p<0.05$. (**G**) Same as E but for SST interneurons. 3-way ANOVA, 20 Hz: Stimulation x Day interaction: $F(1, 23)=4.749$, $p<0.05$. 40 Hz: Main effect of Pulse Number: $F(2.573, 59.17)=15.99$, $p<0.0001$; Main effect of Day: $F(1, 23)=21.57$, $p<0.0001$; Pulse Number x Day interaction: $F(1.759, 40.46)=10.79$, $p<0.0005$; *$p<0.05$, Day 1 vs. Day 22; n=12-13. (H) Same as F but for SST interneurons. Fixed effect of trial reported in gray, $p>0.05$.

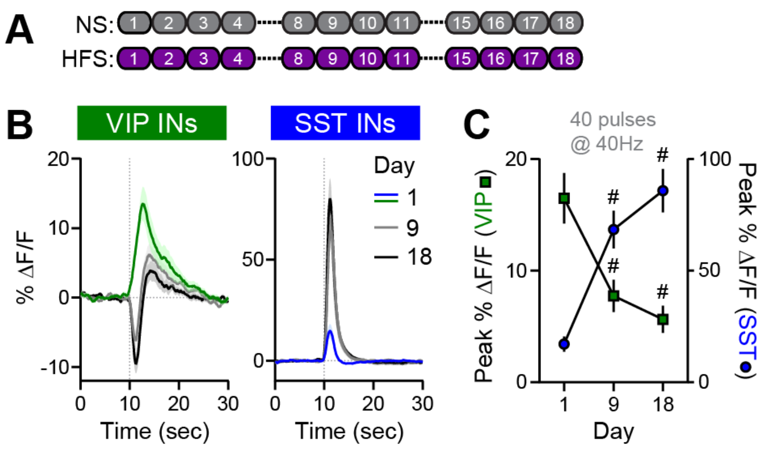


**Figure S5. Repeated HFS alone is sufficient to induce bidirectional plasticity of mPFC VIP and SST interneuron responses.**

(**A**) Schematic of 18-day stimulation protocol consisting of either 12 sessions of no stimulation (NS) or HFS. (**B-C**) Average (**B**) and average peak (**C**) stimulation-evoked $Ca^{2+}$ responses of VIP and SST interneurons in HFS mice on Days 1, 9, and 18 in HFS mice. Stimulation (40 pulses at 40 Hz) initiated at 10-sec timepoint. Mixed-effects model, VIP Main effect of Day: $F(1.237, 14.23)=20.20$, $p<0.0005$; #$p<0.005$, different from Day 1; $n=13$. SST Main effect of Day: $F(1.934, 26.11)=34.13$, $p<0.0001$; #$p<0.0001$, different from Day 1; $n=15$.

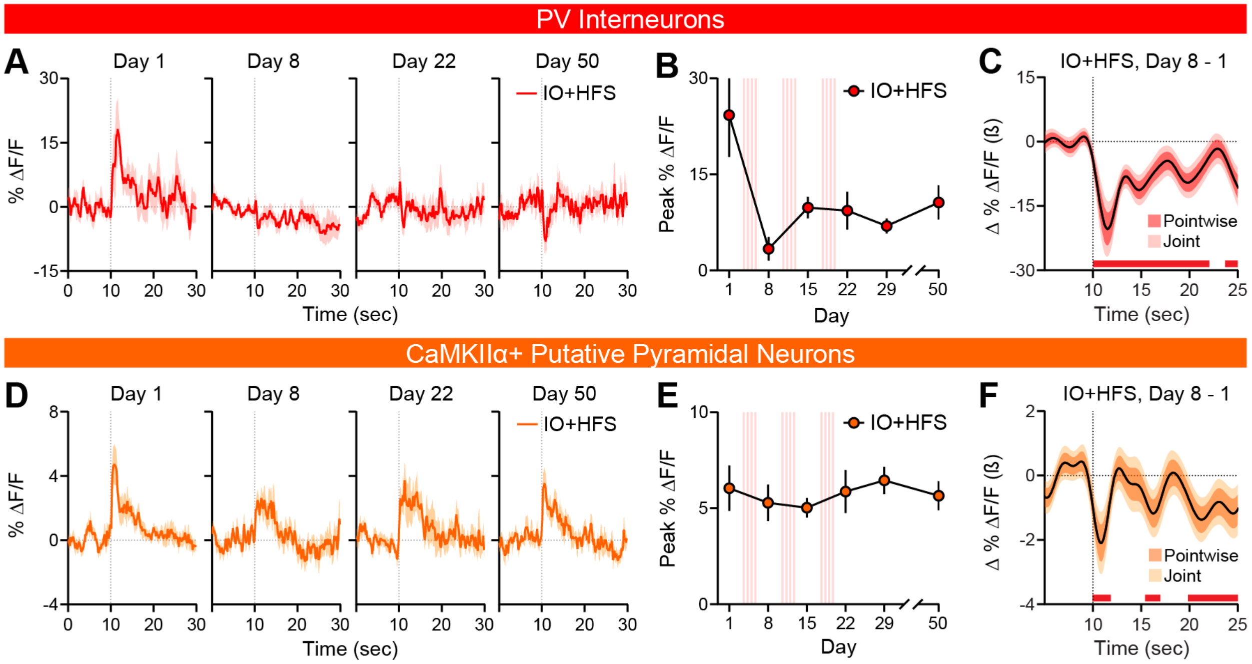


**Figure S6. Repeated vHPC input stimulation produces modest reductions in PV interneuron and pyramidal neuron responses.**

(**A**) Average stimulation-evoked $Ca^{2+}$ responses of mPFC PV interneurons in IO+HFS mice on IO Days 1, 8, 22, and 50. Stimulation (40 pulses at 40 Hz) initiated at 10-sec timepoint. (**B**) Average peak stimulation-evoked $Ca^{2+}$ responses of PV interneurons in IO+HFS mice across all six IO days. Stimulation was 40 pulses at 40 Hz. Repeated measures ANOVA, Main effect of Day: $F(1{,}948, 11.69)=5.472$, $p<0.05$; n=7. (**C**) Functional linear mixed modeling (FLMM) of Day covariate effects on mouse-level PV interneuron $Ca^{2+}$ signals in IO+HFS mice. Functional coefficient estimates of the Day covariate at each timepoint in 30-sec stimulation trials, showing differences in average $Ca^{2+}$ signals in Day 8 relative to Day 1 in IO+HFS mice. Red bars denote post-stimulation timepoints of significant reduction (Joint CIs do not contain Y=0). (**D**) Same as **A** but for CaMKIIα+ putative pyramidal neurons. n=10. (**E**) Same as **B** but for CaMKIIα+ putative pyramidal neurons. (**F**) Same as **C** but for CaMKIIα+ putative pyramidal neurons.

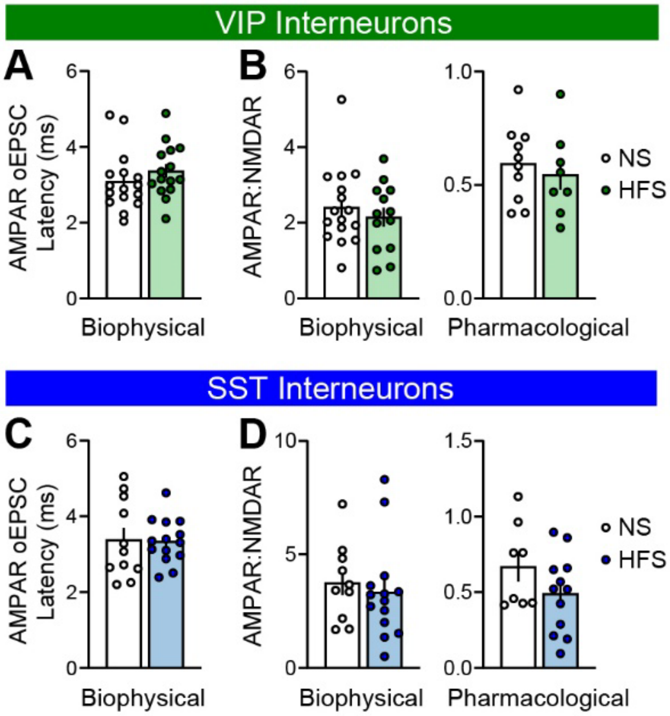


**Figure S7. Additional electrophysiological characterization of monosynaptic vHPC input onto mPFC VIP and SST interneurons.**

(**A**) Latency of biophysically isolated AMPAR-mediated monosynaptic oEPSCs in VIP interneurons of NS and HFS mice (n=15-16 cells). (**B**) Biophysically (left) and pharmacologically (right) isolated AMPAR:NMDAR ratios in VIP interneurons of NS and HFS mice (n=8-16 cells). (**C**) Latency of biophysically isolated AMPAR-mediated monosynaptic oEPSCs in SST interneurons of NS and HFS mice (n=11-14 cells). (**D**) Biophysically (left) and pharmacologically (right) isolated AMPAR:NMDAR ratios in SST interneurons of NS and HFS (n=8-14 cells).

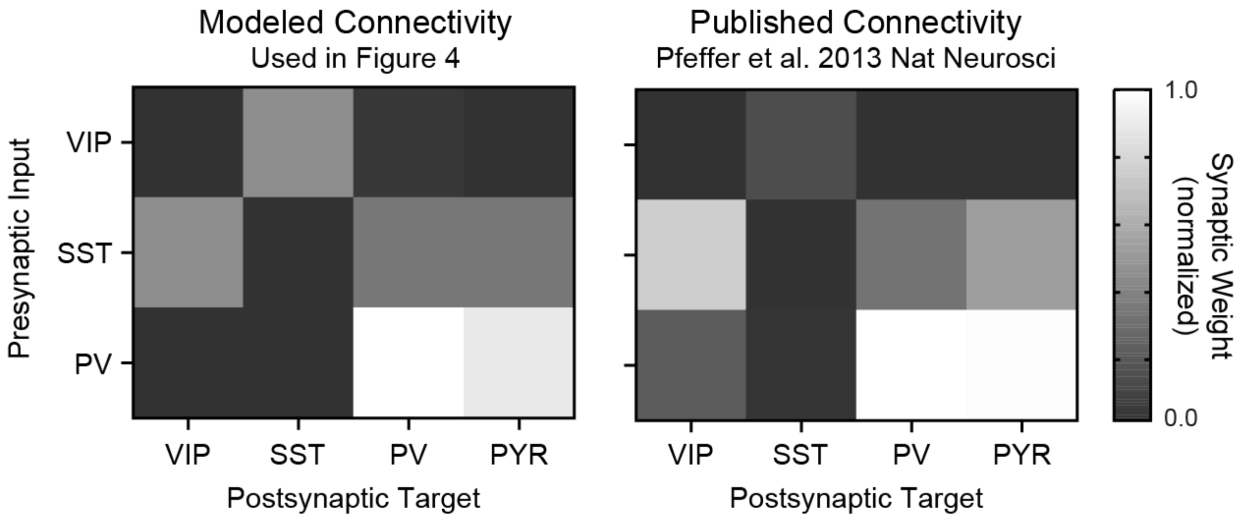


**Figure S8. Cortical microcircuit connectivity implemented in the computational model.**

Heatmaps showing normalized synaptic weights between the four neuronal populations incorporated into the computational model used in Figure 4. The modelled connectivity matrix was based on, and closely approximates, the experimentally measured connectivity described by Pfeffer et al., 2013. Rows denote presynaptic neuronal populations and columns denote postsynaptic targets. Synaptic weights were normalized to the strongest connection in the network.

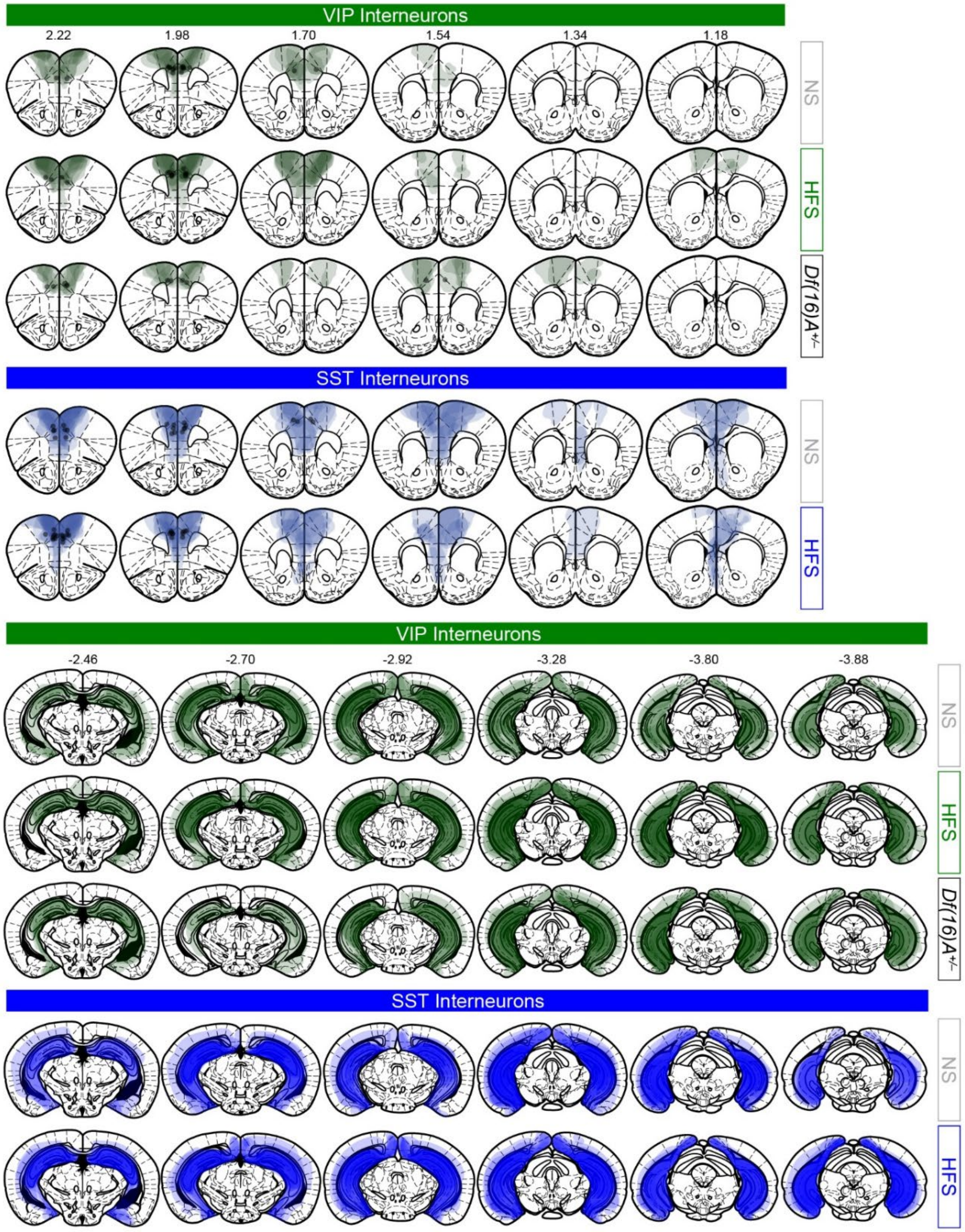


**Figure S9. Histology of ChrimsonR-tdTomato and GCaMP6f expression in mPFC and vHPC and optical fiber placement in mPFC for SWM experiments.**

AAV.CaMKIIα::ChrimsonR-tdTomato expression in vHPC and AAV.Syn::FLEX.GCaMP6f in mPFC VIP and SST interneurons in NS, HFS, and *Df(16)A*$^{+/–}$ mice used for the SWM experiment. Gray circles indicate optical fiber placement.

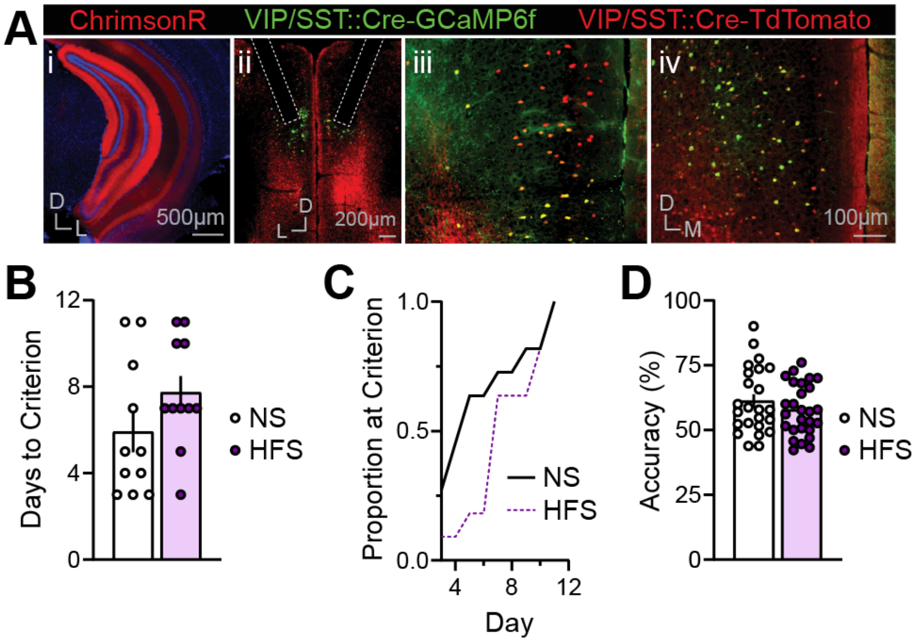


**Figure S10. Prior vHPC input stimulation does not significantly alter overall SWM task acquisition.**

(**A**) Representative viral ChrimsonR-tdTomato expression (red) in vHPC (i) and vHPC terminals in mPFC (ii-iv); genetic tdTomato expression in VIP or SST interneurons (red cell bodies) and viral GCaMP6f expression in VIP or SST interneurons (green; ii-iv). Optic fiber placement denoted by dashed lines. DAPI staining (blue) shown in vHPC (i). (**B**) Days to reach criterion performance (3 consecutive days ≥70% accuracy) in the SWM task in NS and HFS mice (n=11,11). (**C**) Cumulative distributions of NS and HFS mice that reached criterion across training days. (**D**) Average accuracy across training in NS and HFS mice (n=24-27).

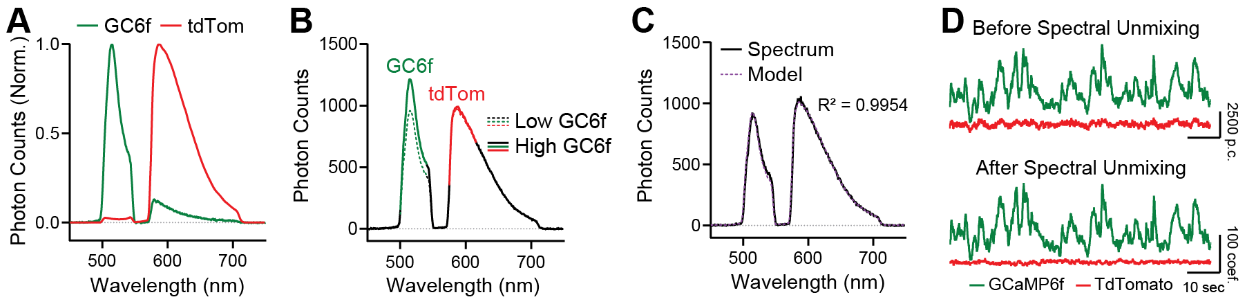


**Figure S11. Spectral unmixing validation.**

(**A**) Normalized photon counts of GCaMP6f and tdTomato reference spectra recorded from mice solely expressing GCaMP6f or tdTomato. (**B**) Photon counts spanning the mixed GCaMP6f-tdTomato spectrum of a representative SST::Cre mouse at two experimental timepoints. The timepoints correspond to high- (solid lines) and low-points (dashed lines) in the GCaMP6f timeseries to show the dynamics changes to the GCaMP portion of the spectrum and relative stability of the tdTomato portion. (**C**) Photon counts of a single representative measured spectrum overlayed with its modeled spectrum generated by the linear unmixing algorithm. (**D**) Representative GCaMP6f and tdTomato traces showing photon counts (before spectral unmixing) and coefficient (after spectral unmixing) of the two signals across time.

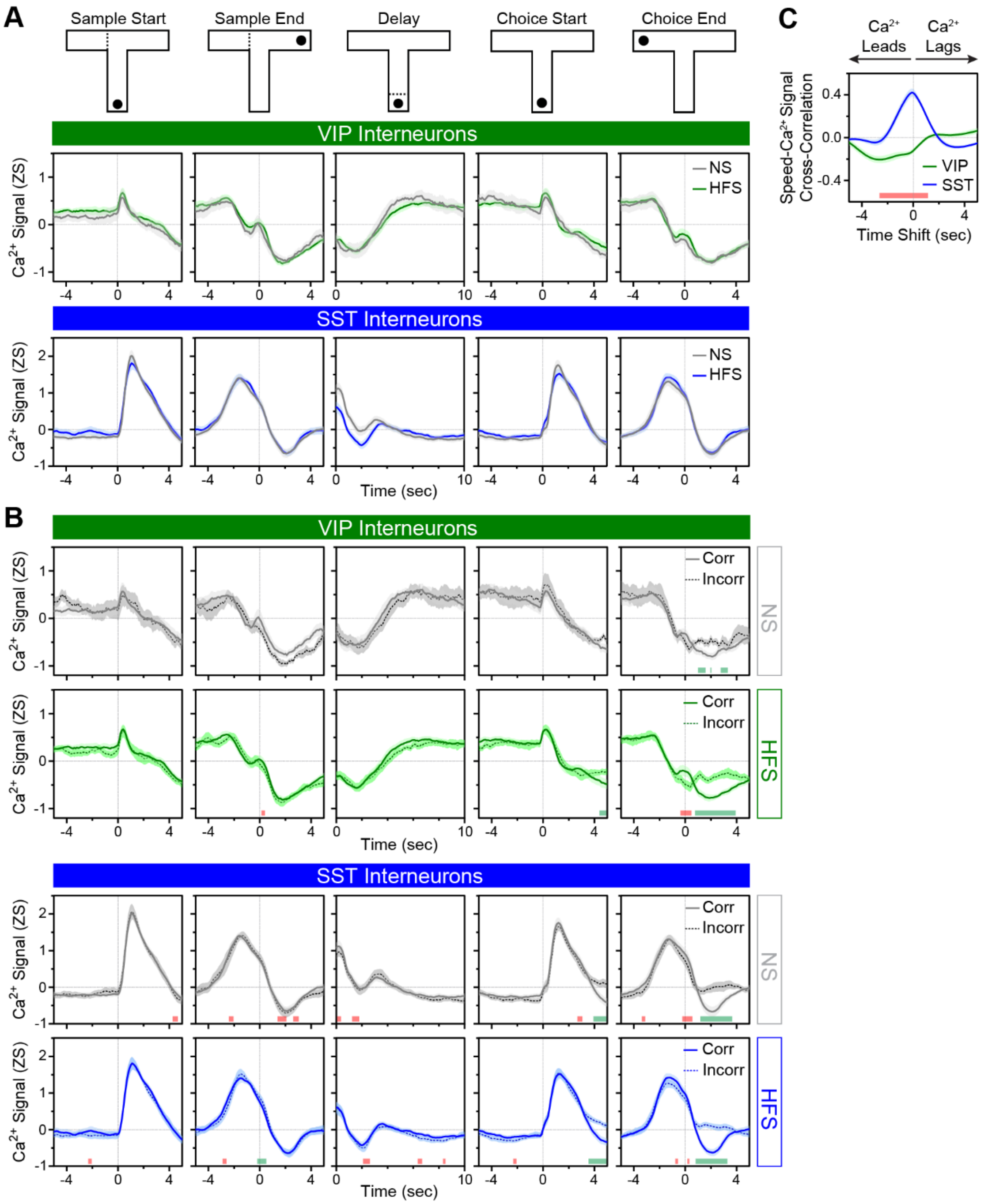


Figure S12. Average task-related activity and speed–activity relationships of mPFC VIP and SST interneurons across all SWM training days.

(**A**) Average Z-scored $Ca^{2+}$ signals of VIP and SST interneurons from NS and HFS mice from correct trials across all training days and aligned to discrete SWM task epochs. n=10-13 VIP; n=14-15 SST. (**B**) Same as A but plotting correct and incorrect trials from all training days. Green and red bars denote timepoints of significant enhancement and reduction in incorrect relative to correct trials, respectively, using functional linear mixed modelling. (**C**) Average cross-correlation of speed and VIP (green) and SST interneurons (blue) or $Ca^{2+}$ signals across whole trials of the SWM task. Red bar denotes timepoints of significant difference between VIP and SST interneuron speed-$Ca^{2+}$ cross-correlations using functional linear mixed modeling. n=10-14.

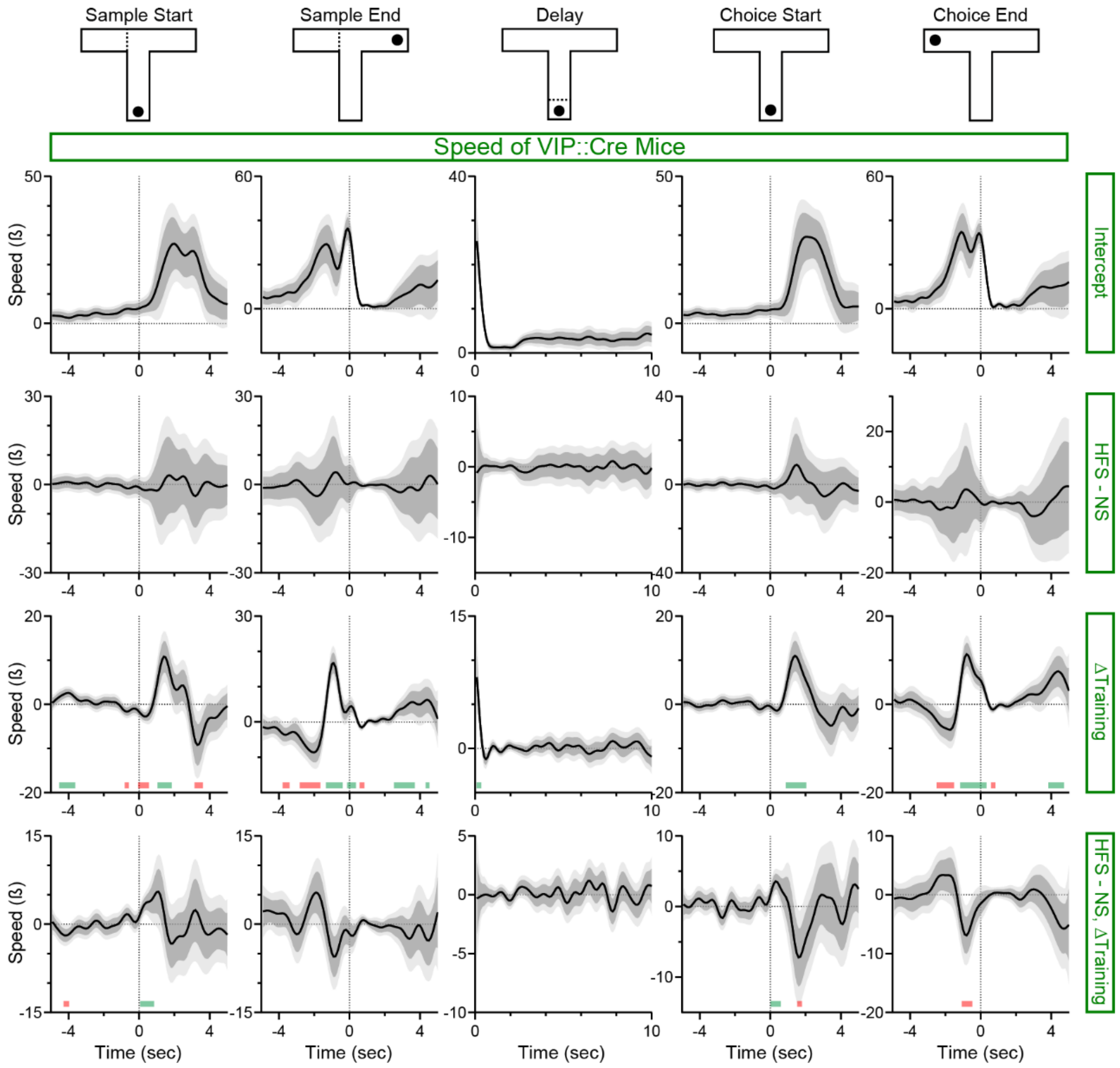


**Figure S13. Functional linear mixed-effects analysis of HFS, training, and HFS × training effects on speed across SWM task epochs in VIP::Cre mice.**

Functional intercept estimates and coefficient estimates of the Stimulation, Training, and the interaction between Stimulation and Training covariate effects on speed in VIP::Cre mice around five SWM task events. Green and red bars denote timepoints of significant enhancement and reduction in coefficient values, respectively.

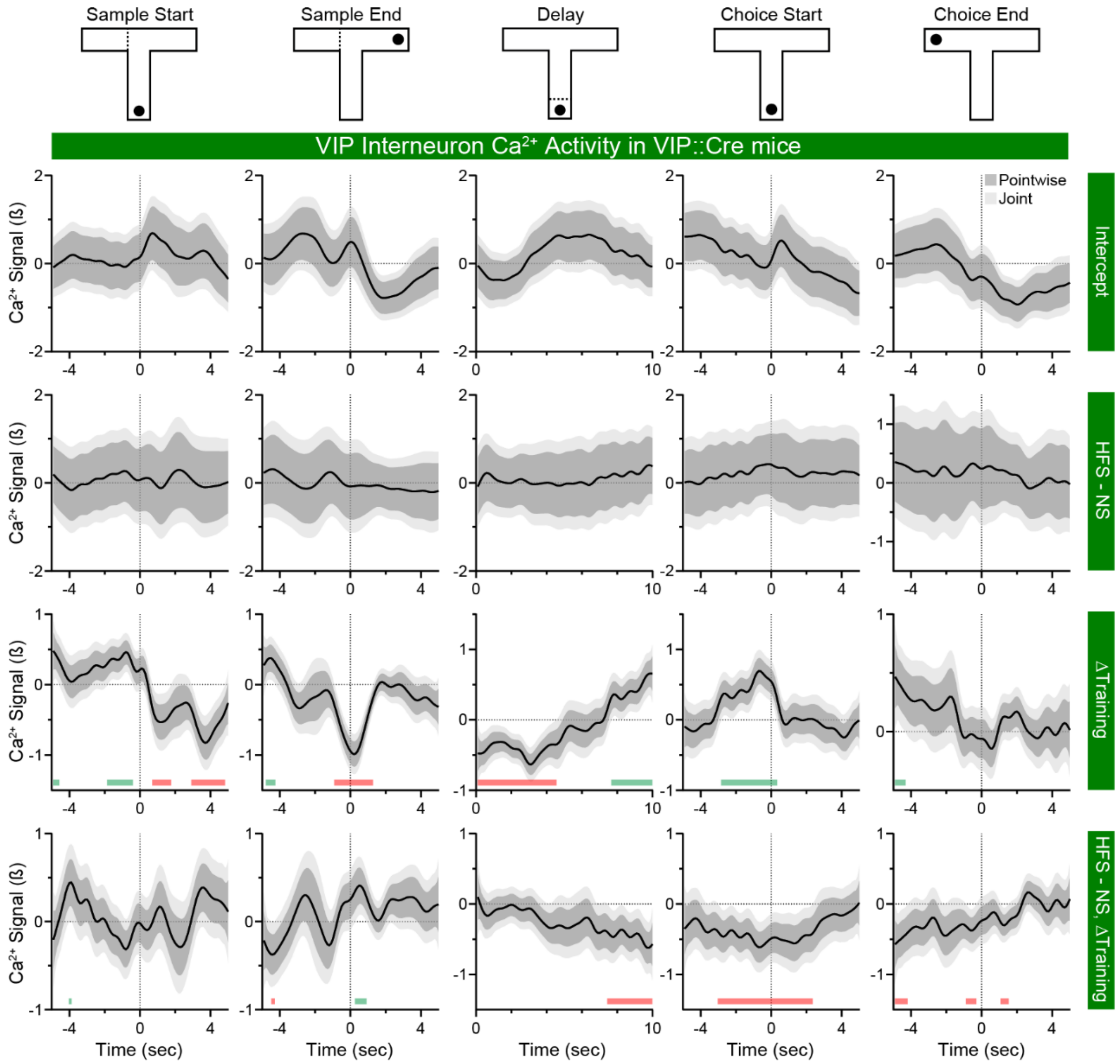


**Figure S14. Functional linear mixed-effects analysis of HFS, training, and HFS × training effects on VIP interneuron $Ca^{2+}$ activity across SWM task epochs.**

Functional intercept estimates and coefficient estimates of the Stimulation, Training, and the interaction between Stimulation and Training covariate effects on VIP interneuron $Ca^{2+}$ signals around five SWM task events. Green and red bars denote timepoints of significant enhancement and reduction in coefficient values, respectively.

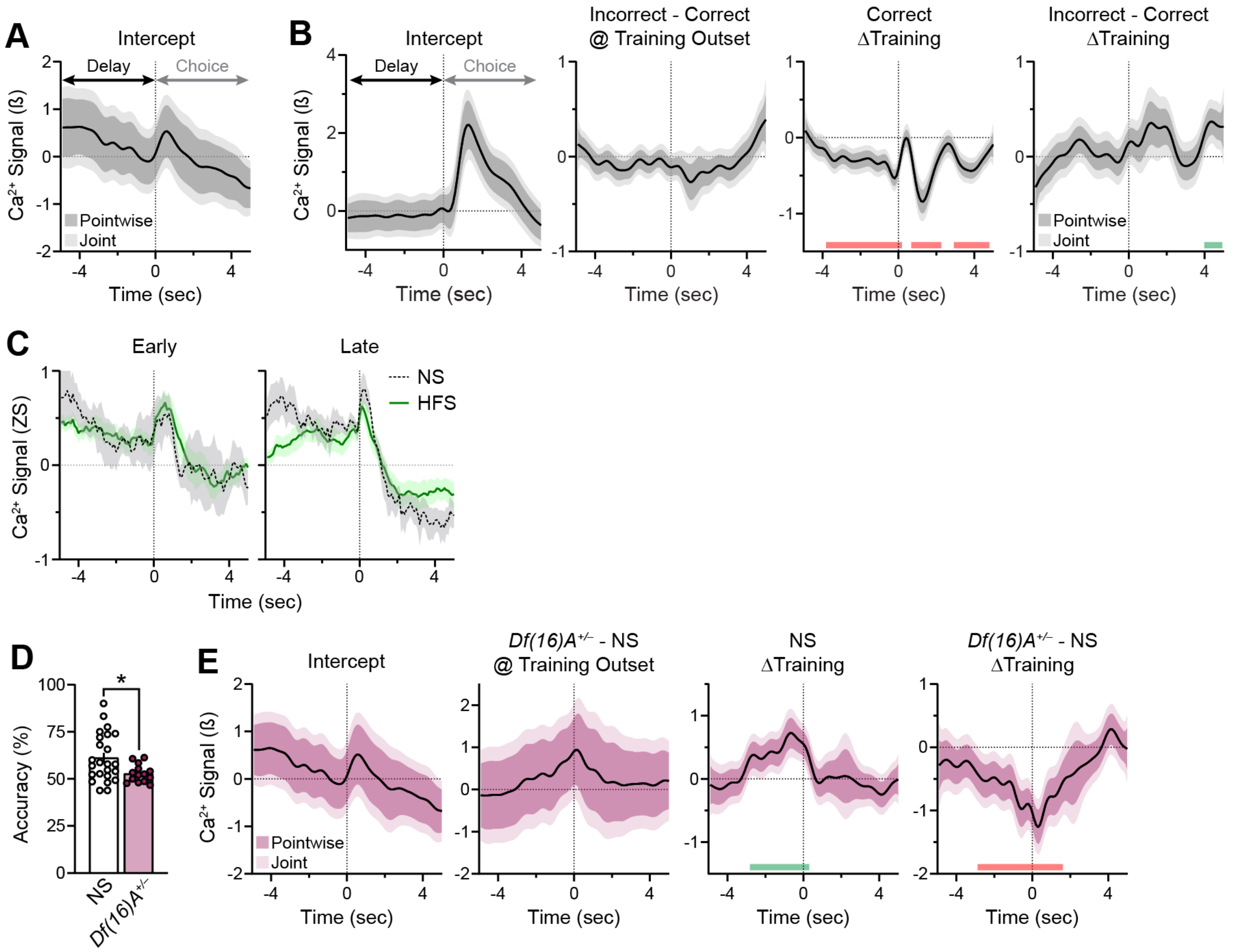


**Figure S15. Additional measures of delay epoch-related VIP interneuron $Ca^{2+}$ activity and their modulation by prior vHPC input stimulation and *Df(16)A*$^{+/–}$ mutation.**

(**A**) Functional intercept estimates of the Outcome, Training, and interaction between Outcome and Training covariates effects on trial-level VIP interneuron $Ca^{2+}$ signals during the Delay-to-Choice transition (See Figure **6**B). **(B)** Functional linear mixed modeling (FLMM) of Outcome and Training covariate effects on trial-level **SST** interneuron $Ca^{2+}$ signals in NS mice during the Delay-to-Choice transition. **Panel 1: Same as A but for SST interneurons. Panel 2**: Functional coefficient estimates of the Outcome covariate at each timepoint in incorrect trials, showing **no** difference**s** in average **SST interneuron** $Ca^{2+}$ signals in incorrect relative to correct trials at the onset of training. **Panel 3**: Functional coefficient estimates of the Training covariate in correct trials, showing changes in average **SST interneuron** $Ca^{2+}$ signals across training. **Panel 4**: Functional coefficient estimates of the interaction between the Training and Outcome covariates, showing **that** changes in average **SST interneuron** $Ca^{2+}$ signals across training **do not** differ in incorrect relative to correct trials. Green and red bars denote timepoints of significant (Joint CIs do not contain Y=0) enhancement and reduction, respectively. (C) Average Z-scored $Ca^{2+}$ signals of VIP interneurons during the Delay-to-Choice transition for incorrect trials in NS and HFS mice in Early and Late stages of training. (D) Average accuracy (%) across training in NS and *Df(16)A*$^{+/-}$ mice. * unpaired t-test: t(36)=2.36, p<0.05; n=24,14. (E) Functional intercept estimates and coefficient estimates of the Genotype, Training, and interaction between Training and Genotype covariates effects on trial-level VIP interneuron $Ca^{2+}$ signals during the Delay-to-Choice transition (See Figure **6**G). Green and red bars denote timepoints of significant enhancement and reduction, respectively.
